# Targeted ubiquitination of Na_V_1.8 reduces sensory neuronal excitability

**DOI:** 10.1101/2025.02.04.636451

**Authors:** Sidharth Tyagi, Mohammad-Reza Ghovanloo, Matthew Alsaloum, Philip Effraim, Grant P. Higerd-Rusli, Fadia Dib-Hajj, Peng Zhao, Shujun Liu, Stephen G. Waxman, Sulayman D. Dib-Hajj

**Author notes:** These authors share senior and last authorship (Stephen G. Waxman, Sulayman D. Dib-Hajj). Corresponding Authors Sidharth Tyagi, PhD, Stephen G. Waxman, MD, PhD Sulayman D. Dib-Hajj, PhD, Neuroscience and Regeneration Research Center, VAMC, 950 Campbell Avenue, Bldg. 34, West Haven, CT 06516, or or.

## Abstract

Chronic pain and addiction are a significant global health challenge. Voltage-gated sodium channel Na_V_1.8, a pivotal driver of pain signaling, is a clinically validated target for the development of novel, non-addictive pain therapeutics. Small molecule inhibitors against Na_V_1.8 have shown promise in acute pain indications, but large clinical effect sizes have not yet been demonstrated and efficacy in chronic pain indications are lacking.

An alternative strategy to target Na_V_1.8 channels for analgesia is to reduce the number of channels that are present on nociceptor membranes. We generated a therapeutic heterobifunctional protein, named UbiquiNa_V_, that contains a Na_V_1.8-selective binding module and the catalytic subunit of the NEDD4 E3 Ubiquitin ligase. We show that UbiquiNav significantly reduces channel expression in the plasma membrane and reduces Na_V_1.8 currents in rodent sensory neurons. We demonstrate that UbiquiNa_V_ is selective for Na_V_1.8 over other Na_V_ isoforms and other components of the sensory neuronal electrogenisome. We then show that UbiquiNa_V_ normalizes the distribution of Na_V_1.8 protein to distal axons, and that UbiquiNa_V_ normalizes the neuronal hyperexcitability in *in vitro* models of inflammatory and chemotherapy-induced neuropathic pain. Our results serve as a blueprint for the design of therapeutics that leverage the selective ubiquitination of Na_V_1.8 channels for analgesia.

## Introduction

Chronic pain is a massive global health burden, affecting more than 30% of the world population, and costing healthcare systems hundreds of billions of dollars per year^1–4^. Therapeutic options for pain are limited and have a spectrum of adverse effects. Treatments for pain that are specific, effective, and non-addictive are urgently needed^5,6^.

The physiological basis of most human pain sensation is the firing of action potentials initiated via sodium flux through voltage-gated sodium (Na_V_) channels, and their subsequent propagation in axons of dorsal root ganglion (DRG) sensory neurons^7^. The Na_V_1.8 isoform is the principal driver of the action potential spike in nociceptors - sensory neurons that respond to noxious stimuli^7,8^. Na_V_1.8 is unique because it is active at much more depolarized membrane potentials compared to other members of the Na_V_ channel family. Additionally, Na_V_1.8 channels recover rapidly after inactivation^9,10^. These characteristics enable Na_V_1.8 channels to carry the majority of inward currents in the rising phase of the nociceptor action potential and to drive repetitive firing behaviors in these sensory neurons^11–14^. Blunting the activity of Na_V_1.8 channels has the potential to reduce nociceptor firing and ameliorate pain^15^. Since Na_V_1.8 channels are present predominantly in peripheral sensory neurons, agents that target Na_V_1.8 can provide analgesia in a non-addictive manner without central side-effects.

To date, the primary strategy for targeting Na_V_ channels for pain treatment has been to directly inhibit channel conductance using small molecule agents^5,6,16^. One such agent, VX-548 (now called Suzetrigine or Journavx), that targets Na_V_1.8 channels has had recent success in phase II and III clinical trials involving patients in post-surgical acute pain settings ^16,17^ and received FDA approval for moderate-to-severe acute pain^18^. However, the drug has a small effect size (∼1-1.5 points on the NPRS clinical pain scale relative to placebo) and thus it may have limited clinical analgesic benefit^15,19^. Additionally, the clinical benefit of the drug, compared to placebo, in chronic pain conditions has not yet been shown^20^. An alternative and potentially more efficacious strategy than direct channel blockade might be to reduce the number of Na_V_1.8 channels that are present along the entire neuronal surface.

A strategy for unlocking this mechanism is to drive internalization and degradation of Na_V_1.8 channels by targeted ubiquitination. Indeed, targeted degradation strategies (canonically using small molecules, e.g., <u>Pro</u>teolysis-<u>Ta</u>rgeting <u>C</u>himeras, PROTACs) have been applied to a variety of intracellular proteins^21^ as well as membrane proteins^22–25^. The general schema is to develop an agent that a) binds selectively to an intracellular epitope on the target of interest, and b) triggers ubiquitination and subsequent degradation of the protein. Though most targeted protein degradation strategies involve small molecules, peptide-based degraders are an emerging technology^26^. We hypothesized that a similar strategy that targets Na_V_ channels preferentially expressed in nociceptors would likely be an effective analgesic.

Here, we achieve isoform-specific targeted reduction of surface Na_V_1.8 channels by making use of a known selective intracellular binder of this channel. The sodium channel and clathrin linker 1 (SCLT1, previously called CAP-1A) binds selectively to the cytoplasmic C-terminus of Na_V_1.8 compared to other Na_V_ channels and specifically reduces Na_V_1.8 current density in native DRG neurons^27^. A module of the SCLT1 protein with homology to myosin-tail domain (MTD) is both necessary and sufficient for the interaction of the protein with Na_V_1.8. In this study, we have linked the catalytic module of a ubiquitin ligase to the Sclt1-MTD and show that this heterobifunctional engineered polypeptide, named UbiquiNa_V_, results in the targeted reduction of Na_V_1.8 at the cell surface and decreases neuronal excitability in an isoform selective manner. We demonstrate this decreased excitability in multiple *in vitro* models of pain, including inflammatory hyperexcitability and chemotherapy-induced peripheral neuropathy. Overall, this work reveals that targeted reduction of sodium channels at the cell surface represents a promising new strategy for chronic pain therapy.

## Results

### Development of a bifunctional protein that binds Na_V_1.8

A heterobifunctional protein that enables targeted ubiquitin-dependent internalization of a membrane protein requires a selective binding module and a ubiquitinating module. We have previously shown that the protein Sclt1 binds the C-terminus of Na_V_1.8 selectively over all other Na_V_ isoforms and that a module of the Sclt1 protein that shares homology to the Myosin Tail Domain (MTD) is necessary and sufficient for this binding^27^. We first confirmed that overexpressing Sclt1 on its own reduces endogenous Na_V_1.8 current density in rat pup DRG neurons (Sclt1: 123.3 ± 22.5 pA/pF; GFP: 288.8 ± 37.5 pA/pF, p<0.01; **Figure 1a-c**) and overexpressing the MTD alone does not produce a significant change in the current density (MTD: 267.7 ± 63.8; GFP: 191.4 ± 27.47 pA/pF, p>0.05; **Figure 1d-f**).

**Figure 1.**
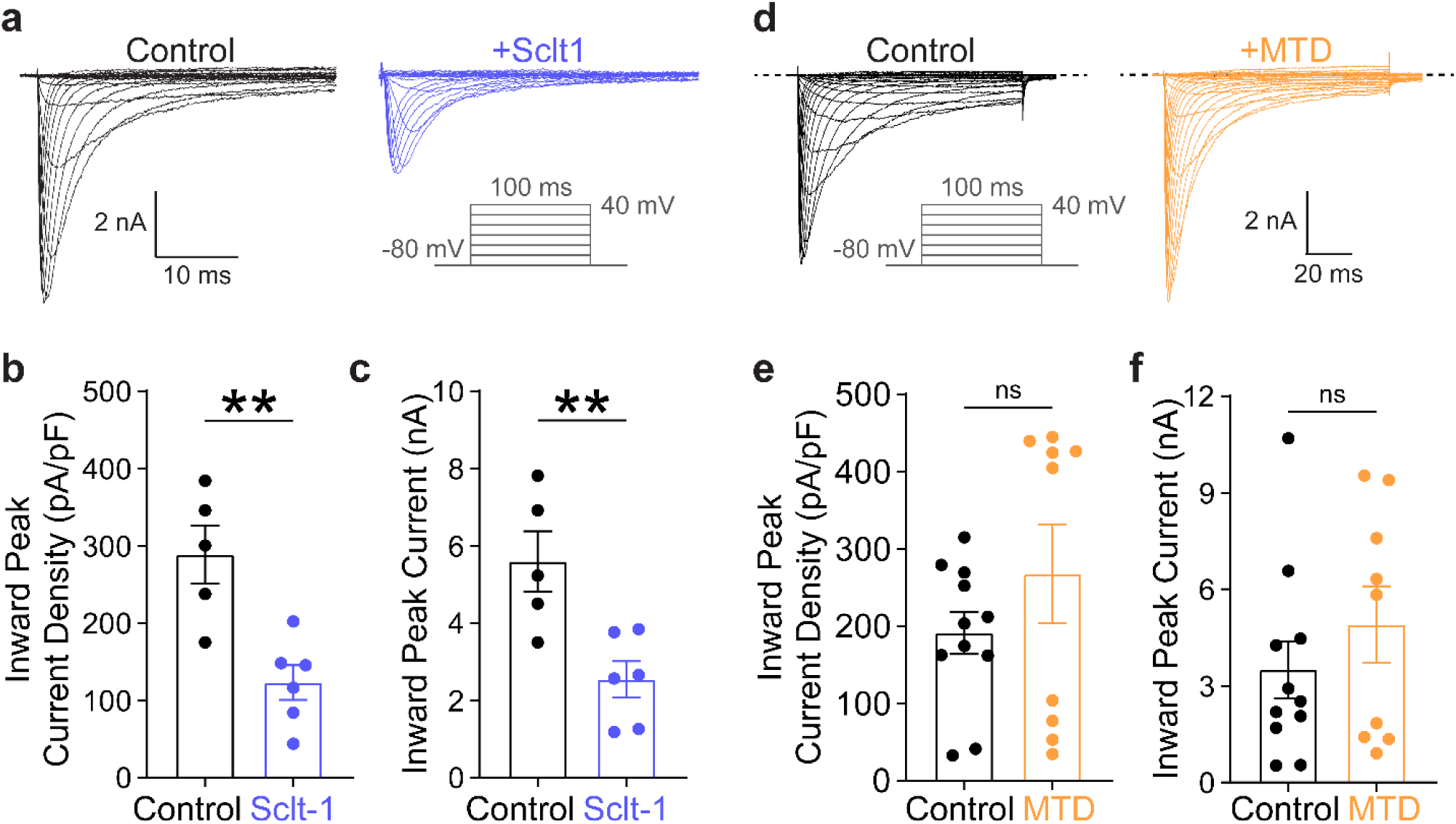
The Sclt-1-MTD, a putative warhead for Na_V_1.8 channels, does not affect Na_V_1.8 currents. (a) Representative whole-cell voltage clamp recordings of DRG neurons from 2-4 d old rat pups transfected with plasmids encoding Sclt1-2A-eGFP or eGFP-control. Inset shows the recording protocol of endogenous TTX-R Na_V_1.8 currents in the presence of 1μM TTX to block endogenous TTX-S currents. (b) Inward peak Na^+^ current density through endogenous Na_V_1.8 channels is significantly reduced in neurons expressing Sclt1-2A-eGFP (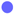; n=6), compared to eGFP-control (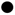; n=5). (c) Inward peak Na^+^ current through endogenous Na_V_1.8 channels is significantly reduced in neurons expressing Sclt1-2A-eGFP (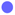; n=6), compared to eGFP-control (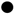; n=5). (d) Representative whole-cell voltage clamp recordings of DRG neurons from 2-4 d old rat pups transfected with plasmids encoding mRuby2-P2A-MTD or mRuby2-control. (e) Inward peak Na^+^ current density through endogenous Na_V_1.8 channels in neurons transfected with mRuby2-P2A-MTD (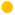; n=9) or mRuby2-control (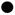; n=11). (f) Inward peak Na^+^ current through endogenous Na_V_1.8 in neurons transfected with mRuby2-P2A-MTD (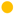; n=9) or mRuby2-control (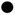; n=11). Bars: Mean ± SEM. ns p> 0.05, ** p<0.01 by the Mann-Whitney U-test.

NEDD4 is a ubiquitin ligase shown to tag Na_V_1.8 and other sodium channel isoforms for internalization^28,29^. Overexpression of NEDD4 in neurons results in a non-selective reduction of multiple types of ion channels and other membrane proteins^30,31^. Previous work has shown that attachment of the NEDD4L (also known as NEDD4-2) catalytic HECT domain to a nanobody specific for neuronal Ca^2+^-channels enables the ubiquitination of these channel targets^23,24^. We attached the HECT domain of NEDD4L to the Sclt1 MTD to generate UbiquiNa_V_ (**Figure 2**). The UbiquiNa_V_ construct contains an eGFP reporter linked in frame by a P2A linker to enable identification of positively transfected cells. We posited that this combination of a warhead and catalytic domain would result in a heterobifunctional protein that binds selectively to Na_V_1.8 channels and results in their ubiquitination and internalization from the plasma membrane.

**Figure 2.**
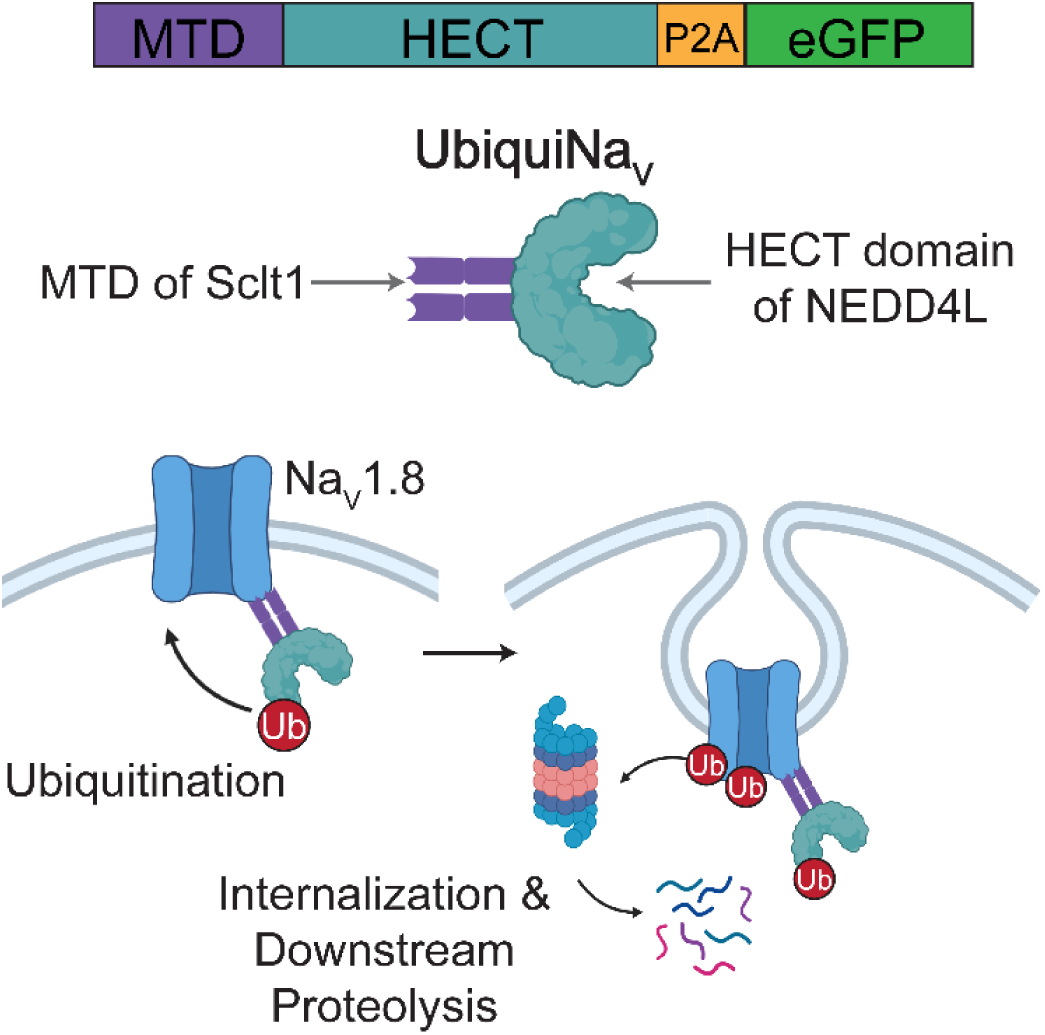
Development of UbiquiNa_V_, a heterobifunctional regulator of Na_V_1.8 channels. UbiquiNa_V_ is composed of two modules separated by a short amino acid linker. The Na_V_1.8 binding module is the MTD of Sclt1, and the catalytic module is the HECT domain of the NEDD4L ubiquitin ligase. An eGFP reporter is fused in frame via a P2A linker.

### UbiquiNa_V_ reduces current density, surface expression, and delivery of Na_V_1.8 channels to distal axons in rat pup DRG neurons

To evaluate the mechanism of action of UbiquiNa_V_ on Na_V_1.8 channels, we transfected P2-4 rat pup DRG neurons with plasmids encoding UbiquiNa_V_, eGFP (to control for the effects of transfection), or a catalytically inactivated UbiquiNa_V_ in which the catalytic activity of the HECT domain is inactivated by a point mutation (C942S^32^) and recorded endogenous Na_V_1.8 currents by whole cell voltage-clamp (**Figure 3a**). DRG neurons expressing UbiquiNa_V_ had significantly decreased Na_V_1.8 current density compared to neurons transfected with an eGFP control (172.5 ± 37.3 vs. 368.4 ± 62.5 pA/pF; p<0.05; **Figure 3b**). Catalytic inactivation of UbiquiNa_V_ ablated these effects on Na_V_1.8 current density (434.6 ± 103.3 pA/pF, p<0.05 vs. UbiquiNa_V_, p>0.05 vs. eGFP-control; **Figure 3b**).

**Figure 3.**
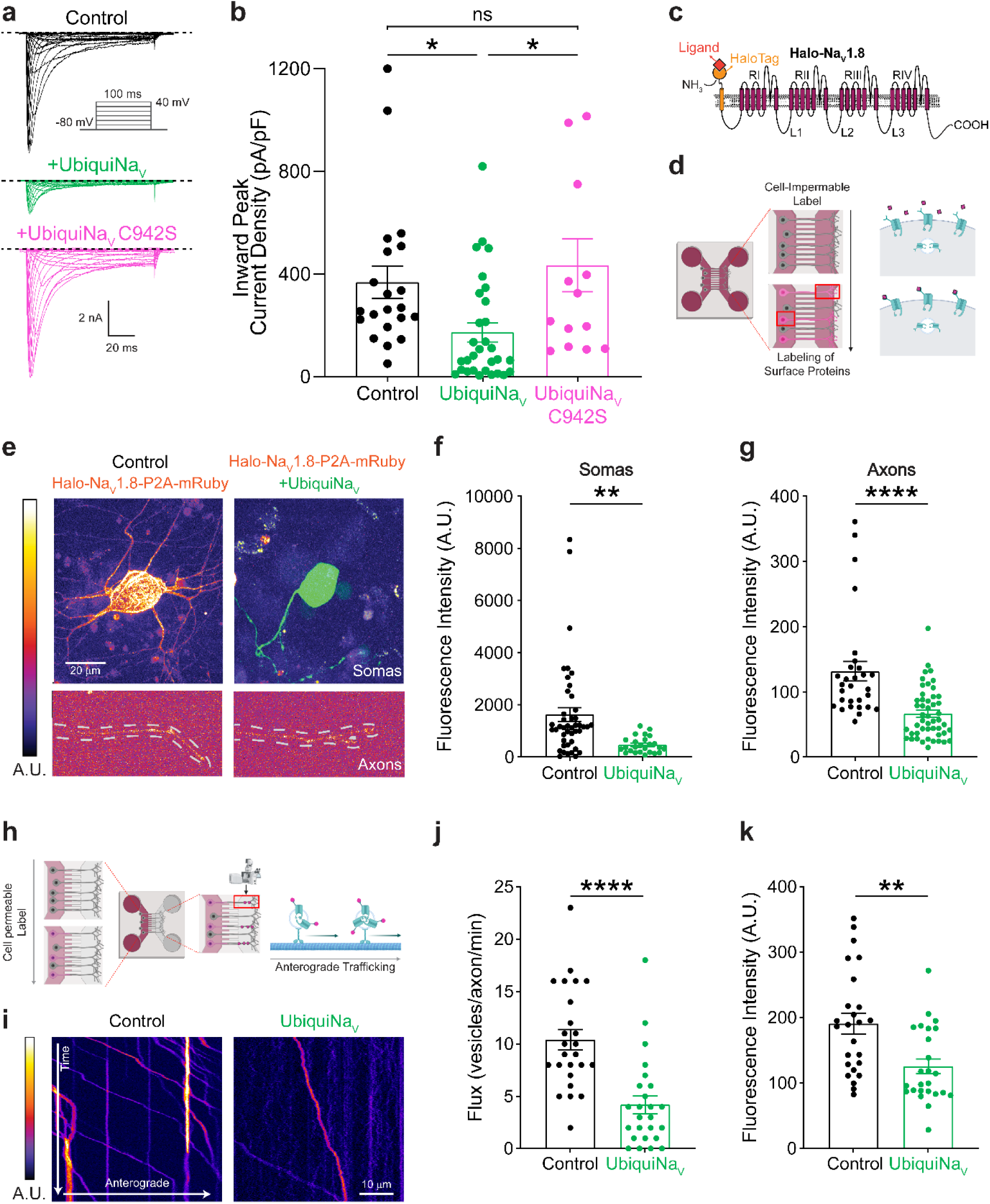
UbiquiNa_V_ decreases Na_V_1.8 current density by way of reducing neuronal surface expression. (a) Representative whole-cell voltage clamp recordings of DRG neurons from 2-4 d old rat pups transfected with plasmids encoding UbiquiNa_V_, UbiquiNa_V_ C942S, or eGFP-control. (b) Inward peak Na^+^ current density through Na_V_1.8 in neurons transfected with UbiquiNa_V_ (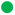; n=30), UbiquiNa_V_ C942S (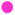; n=14), or eGFP-control (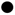; n=21). (c) Schematic depicting the membrane topology of Halo-Na_V_1.8, which contains an extracellular HaloTag enzyme. (d) Schematic of surface labeling experiments. Cell-impermeant JF635i-HaloTag is added to the somatic and axonal chambers of MFCs containing DRG neurons expressing Halo-Na_V_1.8-P2A-mRuby2 and either eGFP-control or UbiquiNa_V_. Following labeling, neurons are fixed with 4% PFA and imaged with confocal microscopy. (e) Representative images of Halo-Na_V_1.8 expression at neuronal surfaces in Halo-Na_V_1.8-expressing neurons transfected with eGFP-control (left panels) or UbiquiNa_V_ (right panels). Top panels – somatic expression. Bottom panels – Axonal expression. mRuby2 fluorescence is not depicted for clarity. (f) Quantification of somatic Halo-Na_V_1.8 surface expression in neurons transfected with Halo-Na_V_1.8-P2A-mRuby2 and eGFP-control (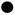; n=25) or UbiquiNa_V_ (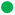; n=44). (g) Quantification of axonal Halo-Na_V_1.8 surface expression in neurons transfected with Halo-Na_V_1.8-P2A-mRuby2 and eGFP-control (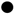; n=29) or UbiquiNa_V_ (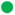; n=49). (h) Schematic of OPAL imaging experiments to evaluate anterograde trafficking of Na_V_1.8. Cell permeable JFX650-HaloTag ligand is added to the somatic chamber of MFCs containing DRG neurons expressing Halo-Na_V_1.8-P2A-mRuby2 and either eGFP-control or UbiquiNa_V_. Following ligand incubation in the somatic chamber, the axonal chamber is imaged by confocal microscopy to reveal anterogradely traveling vesicles carrying labeled Halo-Na_V_1.8. (i) Representative Kymographs from an axon containing anterogradely trafficking vesicles carrying Halo-Na_V_1.8 in neurons transfected with eGFP-control or UbiquiNa_V_. (j) Quantitation of anterograde vesicular flux of Halo-Na_V_1.8 channels in neurons transfected with eGFP-control (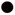; n=25) or UbiquiNa_V_ (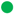; n=25). (k) Quantitation of signal intensity of anterogradely trafficking vesicles carrying Halo-Na_V_1.8 in neurons transfected with eGFP-control (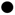; n=25) or UbiquiNa_V_ (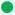; n=25). Bars: Mean ± SEM. ns p> 0.05, * p<0.05, ** p<0.01, **** p<0.0001 by the Mann-Whitney U-test. Multiple comparisons corrected by quantifying the False Discovery Rate and adjusting p-values.

Electrophysiological methods permit the measurement of current density for any given channel type, but the technique only evaluates channel function in the somas of neurons. Sensory afferents *in vivo* are pseudounipolar neurons, each with a long axonal process that extends to the periphery. The distal axonal terminal is the site for initiation of action potentials in response to noxious stimuli^33^. Therefore, a strategy that aims to reduce sodium channel surface expression must accomplish this goal at distal axonal endings. Using a full length Na_V_1.8 construct tagged with the HaloTag enzyme (Halo-Na_V_1.8^34,35^, **Figure 3c**), we evaluated the impact of UbiquiNa_V_ expression on elements of the Na_V_1.8 life cycle in the axons as well as the somas of sensory neurons.

By plating DRG neurons in microfluidic chambers (MFCs), we fluidically isolated axons of DRG neurons from their cell bodies (**Figure 3d**). This approach enabled the evaluation of surface levels of Halo-Na_V_1.8 in both somas and axons^36,37^. We first sought to investigate if Halo-Na_V_1.8 levels at the DRG soma were affected by the presence of UbiquiNa_V_. We applied cell impermeable Halo ligand to the somatic chambers of DRG neurons expressing Halo-Na_V_1.8-P2A-mRuby in glass bottom dishes and then fixed them with PFA (**Figure 3e**). Conjugation to P2A-mRuby was required to identify cells that were positively transfected with Halo-Na_V_1.8 because the concomitant presence of UbiquiNa_V_ would frequently result in very low levels of Halo-Na_V_1.8 at the cell surface. Compared to neurons transfected with eGFP control, neurons transfected with UbiquiNa_V_ exhibited significantly lower levels of Halo-Na_V_1.8 at the somatic surface (461.3 ± 62.0 vs. 1628.0 ± 268.1 A.U., p<0.01; **Figure 3f**). This recapitulates voltage-clamp data which comes from somatic recordings.

Using the same labeling approach but this time in the axonal chamber of MFCs containing DRG neurons, we found that neurons expressing UbiquiNa_V_ had significantly less Halo-Na_V_1.8 channels at distal axonal membranes compared to control (66.6 ± 5.6 vs. 131.5 ± 15.0 A.U., p<0.0001; **Figure 3g**).

This reduction in distal axonal expression of Halo-Na_V_1.8 could be a manifestation of the action of UbiquiNa_V_ at multiple stages of the channel lifecycle. Therefore, we next sought to assess if changes in channel delivery underlie the changes in distal surface expression of Na_V_1.8 induced by UbiquiNa_V_ expression. We investigated how the trafficking of Halo-Na_V_1.8 channels was impacted by the expression of UbiquiNa_V_ using the OPAL imaging technique. By adding cell-permeable fluorophore conjugated Halo ligands to the somatic chamber of MFCs containing neurons transfected with Halo-Na_V_1.8, we can visualize anterogradely trafficking vesicles carrying labeled Halo-Na_V_1.8 channels in axons at a very high signal/noise ratio (**Figure 3h**). We found that the flux of vesicles carrying Halo-Na_V_1.8 is significantly reduced in the presence of UbiquiNa_V_ (4.2 ± 0.9 vs. 10.4 ± 1.0 vesicles/axon/min; **Figure 3i, j**), and that the fluorescence intensity of these vesicles, a measure of the number of labeled channels present, is also reduced (125.7 ± 11.4 vs. 191.0 ± 15.8 A.U.; **Figure 3k**).

To determine if UbiquiNa_V_ travels in vesicles to distal axons, we generated a SNAP-tagged UbiquiNa_V_ construct (**Supplementary** Figure 1a). The SNAPTag functions similarly to HaloTag but binds different cognate ligands which can be co-incubated with Halo ligands. We then transfected Na_V_1.8-null mouse DRG neurons with Halo-Na_V_1.8 and SNAP-UbiquiNa_V_-C942S and evaluated co-trafficking of these proteins with OPAL imaging (**Supplementary** Figure 1b-c). The inactivation of the UbiquiNa_V_ construct was necessary to allow anterograde trafficking of Halo-Na_V_1.8 to travel into the distal axonal compartment since the expression of the active UbiquiNa_V_ significantly reduces the axonal flux (**Figure 3i, j**). We found that >90% of vesicles positive for SNAP-UbiquiNa_V_-C942S are positive for Halo-Na_V_1.8 (**Supplementary** Figure 1d). Additionally, when there were no Na_V_1.8 channels (either endogenous channels or Halo-Na_V_1.8) and Na_V_1.8-null neurons were transfected with the SNAP-UbiquiNa_V_-C942S construct, there were no anterogradely trafficking vesicles carrying SNAP-UbiquiNa_V_-C942S in the axonal chamber (**Supplementary** Figure 1e), whereas robust SNAP-UbiquiNa_V_-C942S could be detected with Halo-Na_V_1.8 co-transfection (**Supplementary** Figure 1f). This data shows that axonal transport of UbiquiNa_V_ is dependent on the presence of Na_V_1.8 channels.

### UbiquiNa_V_ is selective for Na_V_1.8

Na_V_1.8 is an attractive target for pain relief because of its preferential expression in peripheral sensory neurons^7,38,39^. Isoform selectivity of a Na_V_1.8-targeting agent is of paramount importance because other Na_V_ isoforms are expressed in critical electrogenic organs, such as the CNS, autonomic nervous system and cardiac pacemaker cells^40–44^. In peripheral sensory neurons, Na_V_ isoforms can be distinguished by their pharmacologic susceptibility to tetrodotoxin (TTX). Na_V_1.1, Na_V_1.6, and Na_V_1.7 are TTX-Sensitive (TTX-S) while Na_V_1.8 and Na_V_1.9 are TTX-Resistant (TTX-R). These channel isoforms have different biophysical properties, and Na_V_1.8 currents can be isolated by voltage-protocols that inactivate all TTX-S isoforms^45–47^ (**Supplementary** Figure 2a). We found that the expression of UbiquiNa_V_ did not have any effect on TTX-S current density (**Supplementary** Figure 2b-c) in rat pup DRG neurons.

To evaluate the effect of UbiquiNa_V_ on all human Na_V_ isoforms, we designed an automated patch clamp (APC) assay to enable high-throughput, unbiased screening (**Figure 4a**). Here, suspension HEK293 (Expi293F) cells were transfected with plasmids encoding P2A-eGFP-tagged channel constructs and UbiquiNa_V_-P2A-mCherry (or mCherry alone as a control). 48 hours post transfection, we used FACS to isolate the population of cells that expressed fluorescence in both red and green channels, indicating successful co-transfection and expression. We then assayed these cells on the Sophion Qube 384 APC robot. As a positive control, we transfected ND7/23 cells with plasmids encoding eGFP-P2A-Na_V_1.8 and UbiquiNa_V_-P2A-mCherry or mCherry control and proceeded with an identical FACS to APC procedure; ND7/23 cells were used because Na_V_1.8 channels do not produce a robust current in HEK293 cells. UbiquiNa_V_ did not reduce the current density of Na_V_ isoforms 1.1-1.7 (**Figures 4b-h**), nor did it affect the current-voltage (IV) relationships of any of these channels (**Supplementary** Figure 3). However, the reduction in Na_V_1.8 current density expected from UbiquiNa_V_ was recapitulated in this assay (38.9 ± 6.5 vs. 94.1 ± 18.0 pA/pF in UbiquiNa_V_ vs. Control treated cells, respectively; p<0.05; **Figure 4i**). The Na_V_ channel isoform Na_V_1.9 does not express well in heterologous systems^48^. To evaluate the effect of UbiquiNa_V_ on Na_V_1.9, we transfected rat pup SCG neurons (which do not express TTX-R Na_V_ channels natively^49,50^) with plasmids encoding eGFP-2A-hNa_V_1.9 and UbiquiNa_V_-2A-mCherry and assessed current density by manual patch clamp, demonstrating no difference in hNa_V_1.9 current density due to UbiquiNa_V_ (**Supplementary** Figure 4).

**Figure 4.**
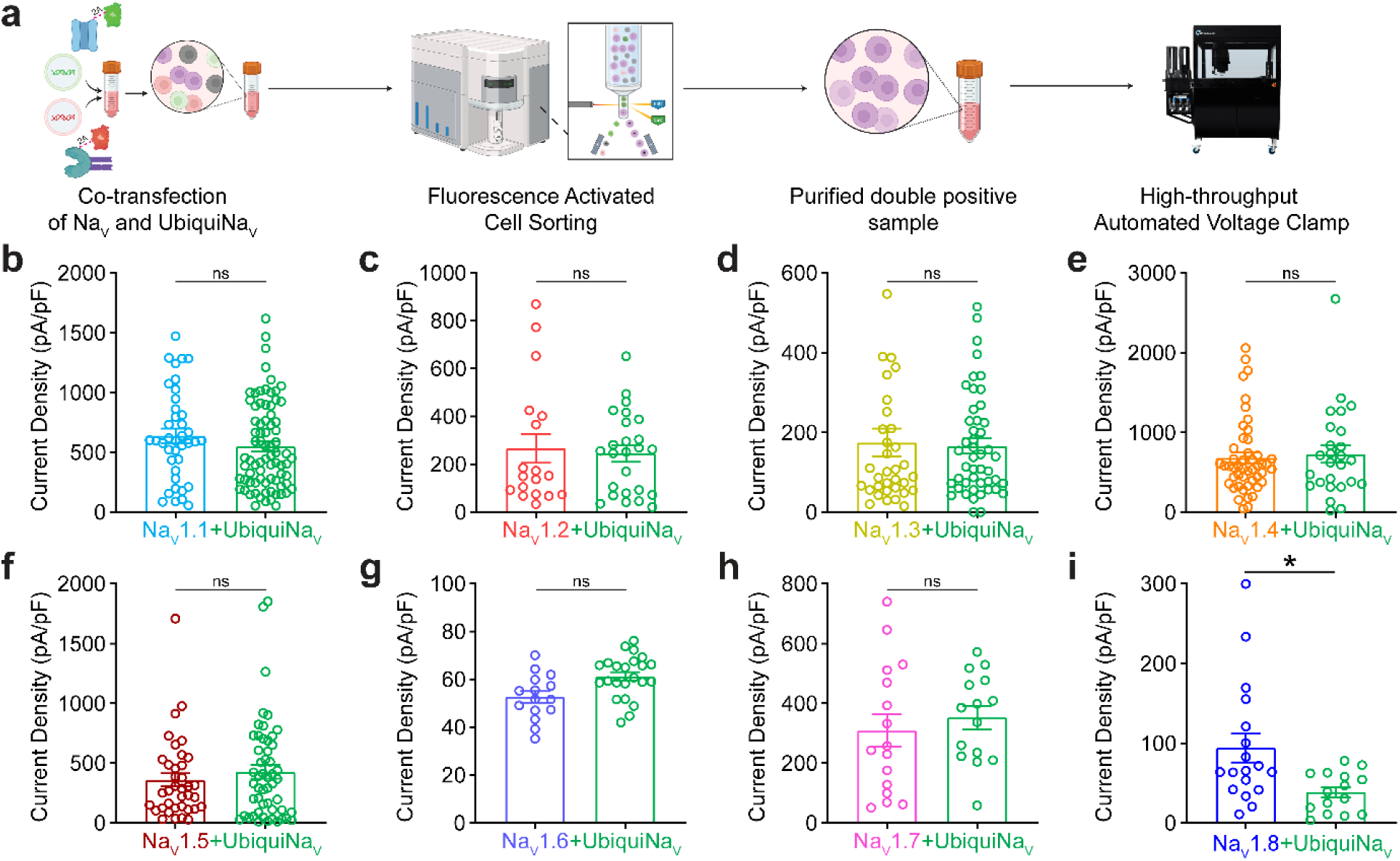
UbiquiNa_V_ is selective for Na_V_1.8. **(a)** Schematic of high-throughput screening assay for the unbiased evaluation of UbiquiNa_V_-mediated effects on human Na_V_ isoforms. Expi293F (suspension) are co-transfected with plasmids encoding eGFP-P2A-Na_V_ channels and either UbiquiNa_V_-P2A-mCherry or mCherry-control. Fluorescence activated cell sorting is conducted to isolate cells expressing both proteins. Whole cell voltage clamp recordings are achieved using the 384-well Sophion Qube Automated Patch Clamp robot. **(b)** Peak inward current densities from automated whole-cell voltage clamp of post-FACS Expi293F cells expressing eGFP-2A-Na_V_1.1 and either mCherry control (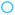; n=39) or UbiquiNa_V_-P2A-mCherry (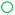; n=77) **(c)** Peak inward current densities from automated whole-cell voltage clamp of post-FACS Expi293F cells expressing eGFP-2A-Na_V_1.2 and either mCherry control (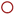; n=18) or UbiquiNa_V_-P2A-mCherry (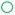; n=25) **(d)** Peak inward current densities from automated whole-cell voltage clamp of post-FACS Expi293F cells expressing eGFP-2A-Na_V_1.3 and either mCherry control (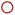; n=34) or UbiquiNa_V_-P2A-mCherry (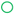; n=45) **(e)** Peak inward current densities from automated whole-cell voltage clamp of post-FACS Expi293F cells expressing eGFP-2A-Na_V_1.4 and either mCherry control (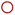; n=48) or UbiquiNa_V_-P2A-mCherry (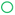; n=25) **(f)** Peak inward current densities from automated whole-cell voltage clamp of post-FACS Expi293F cells expressing eGFP-2A-Na_V_1.5 and either mCherry control (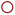; n=25) or UbiquiNa_V_-P2A-mCherry (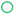; n=26) **(g)** Peak inward current densities from automated whole-cell voltage clamp of post-FACS Expi293F cells expressing eGFP-2A-Na_V_1.6 and either mCherry control (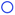; n=15) or UbiquiNa_V_-P2A-mCherry (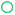; n=23) **(h)** Peak inward current densities from automated whole-cell voltage clamp of post-FACS Expi293F cells expressing eGFP-2A-Na_V_1.7 and either mCherry control (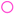; n=16) or UbiquiNa_V_-P2A-mCherry (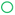; n=15) **(i)** Peak inward current densities from automated whole-cell voltage clamp of post-FACS ND7/23 cells expressing eGFP-2A-Na_V_1.8 and either mCherry control (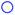; n=18) or UbiquiNa_V_-P2A-mCherry (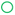; n=15). 1 μM TTX was included in the bath to block endogenous TTX-S channels in ND7/23 cells. Bars: Mean ± SEM. ns p> 0.05, * p<0.05 by the Mann-Whitney U-test (panels b, c, d, e, f, i), or Student’s unpaired t-test (panels g, h).

### UbiquiNa_V_ regulates neuronal excitability in a Na_V_1.8 dependent manner

The sensory neuronal electrogenisome is comprised of a wide variety of voltage-gated ion channels beyond just Na_V_ channels^51–53^. It is unfeasible to individually evaluate the effect of UbiquiNa_V_ on all voltage-gated ion channels that are present in sensory neurons. However, the binding module of UbiquiNa_V_ may bind off-target to other components of the excitability machinery in the cell. To evaluate if this is the case, we employed two parallel approaches to determine if the reduction in neuronal excitability provided by UbiquiNa_V_ is dependent on the presence of Na_V_1.8 channels and the binding of the agent (**Figure 5a**).

**Figure 5.**
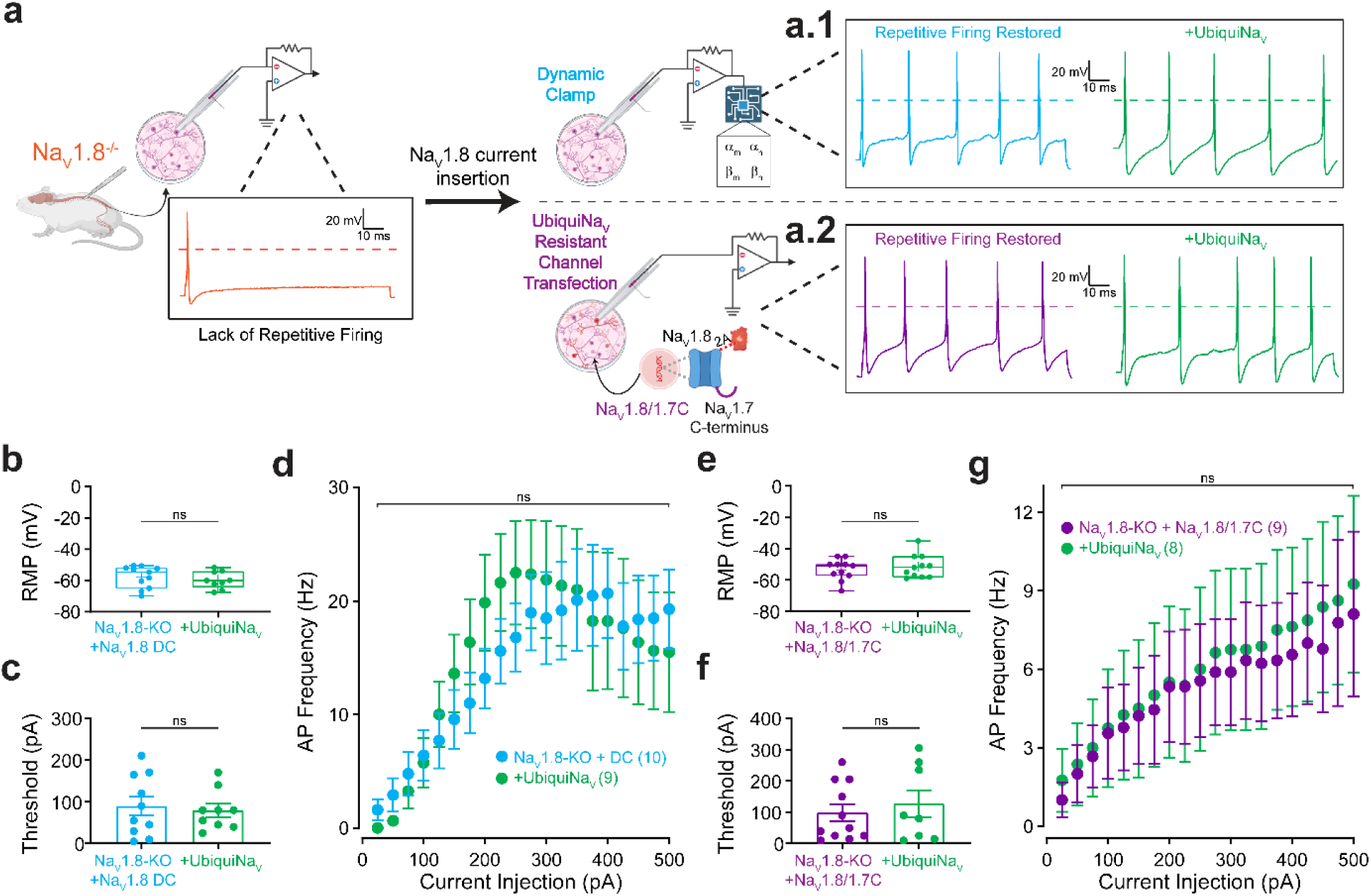
UbiquiNa_V_ does not affect non-Na_V_1.8 components of the neuronal electrogenisome. (a) Schematic of parallel approaches to elucidate the effects of UbiquiNa_V_ expression on the non-Na_V_ electrogenisome. DRG neurons from Na_V_1.8-null mice do not fire repetitive action potentials (representative trace, yellow). Two methods restore repetitive firing in these neurons - dynamic clamp insertion of mathematically modeled Na_V_1.8 conductance (representative blue trace) and transfection of a chimeric Na_V_1.8 channel (containing the Na_V_1.7 c-terminus) that is resistant to UbiquiNa_V_ binding (representative purple trace). Addition of UbiquiNa_V_ in each of these systems does not affect AP morphology (representative green traces). (b) Resting membrane potential recordings from Na_V_1.8-null mouse DRG neurons transfected with either eGFP-control (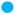) or UbiquiNa_V_ (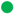) following dynamic clamp addition of Na_V_1.8 current. (c) Action potential firing thresholds recorded from Na_V_1.8-null mouse DRG neurons transfected with either eGFP-control (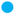) or UbiquiNa_V_ (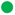) following dynamic clamp addition of Na_V_1.8 current. (d) Repetitive action potential firing in response to gradually increasing current injections (25 pA-500 pA) of 1 s duration in Na_V_1.8-null mouse DRG neurons transfected with either eGFP-control (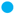) or UbiquiNa_V_ (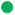) following dynamic clamp addition of Na_V_1.8 current. (e) Resting membrane potential recordings from Na_V_1.8-null mouse DRG neurons transfected with either eGFP-control (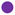) or UbiquiNa_V_ (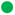) and mCherry-P2A-Na_V_1.8/1.7C. (f) Action potential firing thresholds recorded from Na_V_1.8-null mouse DRG neurons transfected with either eGFP-control (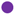) or UbiquiNa_V_ (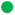) and mCherry-P2A-Na_V_1.8/1.7C. (g) Repetitive action potential firing in response to gradually increasing current injections (25 pA-500 pA) of 1 s duration in Na_V_1.8-null mouse DRG neurons transfected with either eGFP-control (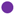) or UbiquiNa_V_ (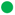) and mCherry-P2A-Na_V_1.8/1.7C. Bars: Mean ± SEM. ns p> 0.05, by Student’s unpaired t-test (panels b, c, e, f), or linear mixed-effects modeling with post-hoc comparison (panels d, g). Multiple comparisons corrected by quantifying the False Discovery Rate and adjusting p-values.

Dynamic clamp is an electrophysiologic method that enables injection of simulated ion channel conductance using a computer-cell interface^54,55^. We first created a computer model of Na_V_1.8 that simulates its physiological activity with high fidelity (**Supplementary** Figure 5a). DRG neurons from Na_V_1.8-null mice demonstrate reduced action potential amplitude and loss of repetitive firing ability (**Figure 5a.1**). We then dynamically injected in physiologic levels of Na_V_1.8-simulated current back into these neurons and found that this addition restores action potential amplitude and repetitive firing behavior (**Supplementary** Figure 5b-c). Then, we compared the action potential firing properties of DRG neurons taken from Na_V_1.8-null mice transfected with UbiquiNa_V_ vs. eGFP-control that had been dynamically injected with Na_V_1.8 current. We found that there was no difference in resting membrane potential (−59.7 ± 1.8 vs. −57.6 ± 2.1 mV, p>0.05; **Figure 5b**), action potential firing threshold (79.4 ± 15.9 vs. 90.0 ± 23.0 pA, p>0.05; **Figure 5c**), or repetitive firing behavior (p >0.05; **Figure 5d**) in DRG neurons transfected with UbiquiNa_V_ vs. eGFP-control in the absence of physical Na_V_1.8 channels.

Though these experiments provided strong evidence to support the idea that UbiquiNa_V_’s effects are dependent on the physical presence of Na_V_1.8 channels, dynamic clamp is an artificial system. To tackle the problem of electrogenic specificity in another way, we leveraged our knowledge of the previously published binding site of the MTD – the C-terminus of Na_V_1.8^27^. We swapped the C-terminus of Na_V_1.8 with the C-terminus of Na_V_1.7 (Na_V_1.8/1.7C), producing a chimeric channel (**Supplementary** Figure 6a) that supported biophysically similar currents to Na_V_1.8 (**Supplementary** Figure 6b) but was resistant to UbiquiNa_V_ activity since the binding site had been altered (**Supplementary** Figure 6c). Transfecting DRG neurons from Na_V_1.8-null mice with this chimeric channel construct restored repetitive firing ability (**Figure 5a.2**). We then assessed if UbiquiNa_V_ had any effects on excitability in Na_V_1.8-null mouse neurons expressing this UbiquiNa_V_-resistant Na_V_1.8/1.7C construct. Recapitulating results of the dynamic clamp experiments, we found that UbiquiNa_V_ did not affect resting membrane potential (−51.5 ± 2.2 vs. −53.4 ± 2.0 mV, p>0.05; **Figure 5e**), action potential firing threshold (127.5 ± 42.5 vs. 97.7 ± 27.5 pA, p>0.05; **Figure 5f**), or repetitive firing behavior (p >0.05; **Figure 5g**) in Na_V_1.8-null mouse DRG neurons transfected with UbiquiNa_V_ vs. eGFP-control along with Na_V_1.8/1.7C.

### Expression of UbiquiNa_V_ normalizes delivery and distribution of Na_V_1.8 channels and neuronal excitability in hyperexcitable rat pup DRG neurons

Our results indicate that UbiquiNa_V_ reduces the levels of Na_V_1.8 channels at the plasma membrane and current density. However, its viability as a potential analgesic depends on its ability to normalize nociceptor hyperexcitability. To evaluate this further, we used current-clamp electrophysiology to investigate the firing properties of rat pup DRG neurons transfected with UbiquiNa_V_ (**Figure 6a**). At baseline, rat pup DRG neurons do not fire trains of action potentials readily^56,57^. However, treatment of cultures with TNF-α rapidly sensitizes neurons and mimics an inflammatory pain state^57–60^. After incubation with 40 ng/mL TNF-α for 24 hours, sensory neurons become hyperexcitable and fire multiple action potentials readily (4.2 ± 1.1 Hz in response to 200 pA; **Figure 6b**). In neurons expressing UbiquiNa_V_, application of TNF-α does not evoke hyperexcitability, and firing frequency in response to current injection was similar to control neurons treated with DMSO vehicle (0.7 ± 0.1 vs. 1.2 ± 0.4 Hz in response to 200 pA; p>0.05, p<0.01 vs. TNF-α) (**Figure 6b**). While TNF-α decreases the threshold for AP firing, neurons that express UbiquiNa_V_ have similar AP thresholds to control neurons **(Figure 6c)**. TNF-α dramatically increases repetitive firing behaviors in response to increasing current injections. UbiquiNa_V_ expression ablates this effect of TNF-α, and neuronal firing remains consistent with un-inflamed firing behavior when UbiquiNa_V_ is on board. (**Figure 6d**)

**Figure 6.**
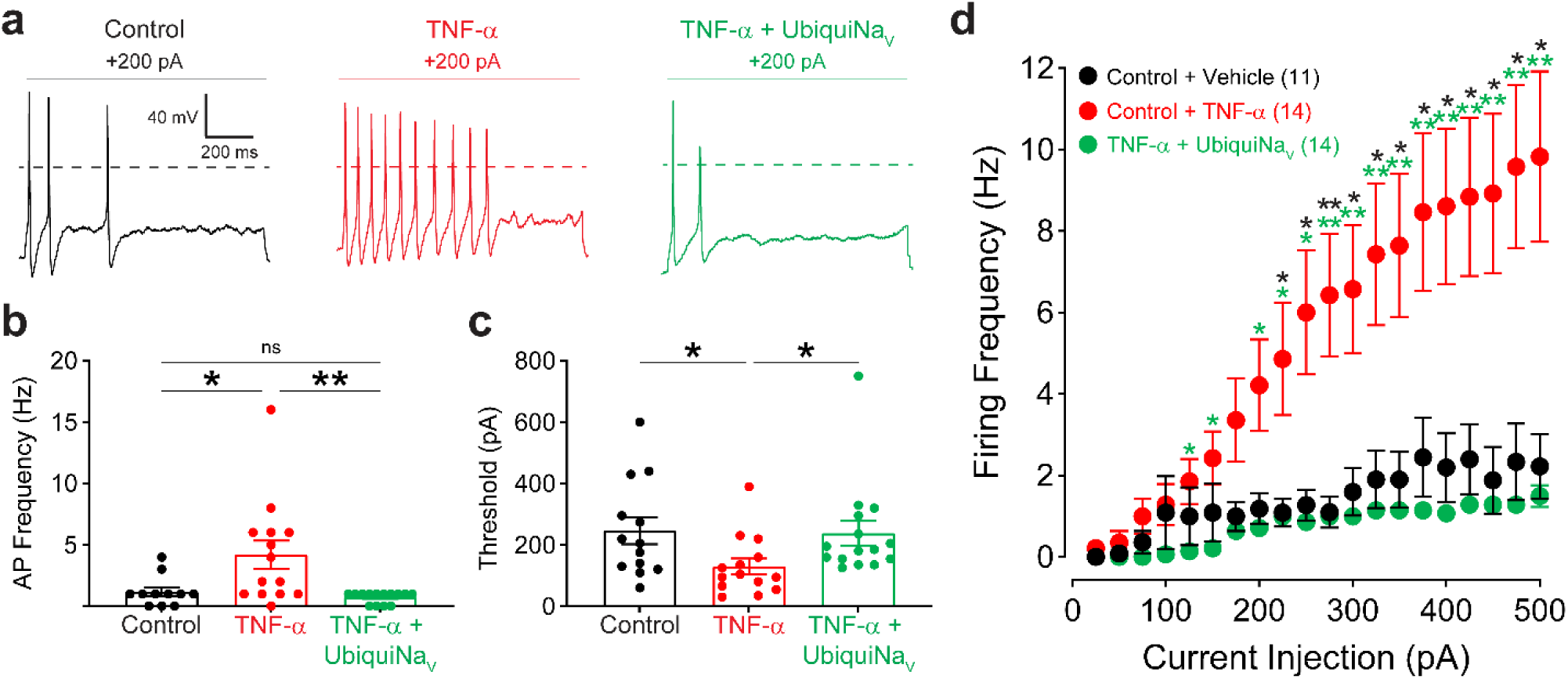
UbiquiNa_V_ normalizes neuronal excitability in a hyperexcitable state. **(a)** Representative whole-cell current clamp recordings of DRG neurons from 2-4 d old rat pups in response to injection of 200 pA of current for 1 s. Control: transfected with eGFP, incubated with 40 ng/mL DMSO. TNF-α: transfected with eGFP, incubated with 40 ng/mL TNF-α for 24 hours. TNF-α + UbiquiNa_V_: transfected with UbiquiNa_V_, incubated with 40 ng/mL TNF-α for 24 hours. Rat pup DRG neurons do not fire readily at baseline. Application of the inflammatory cytokine TNF-α mimics an inflammatory pain state and results in neuronal hyperexcitability. DRG neurons expressing UbiquiNa_V_ do not become hyperexcitable in response to TNF-α exposure. **(b)** Action potentials fired in response to 200 pA current injection in control (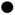; n=11), TNF-α treated (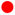; n=14) or UbiquiNa_V_ expressing + TNF-α treated (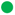; n=14) DRG neurons. **(c)** Action potential firing threshold in control (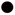; n=11), TNF-α treated (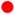; n=14) or UbiquiNa_V_ expressing + TNF-α treated (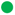; n=14) DRG neurons. **(d)** Repetitive action potential firing in response to gradually increasing current injections (25 pA-500 pA) of 1 s duration in control (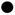; n=11), TNF-α treated (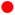; n=14) or UbiquiNa_V_ expressing + TNF-α treated (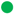; n=14) DRG neurons. Bars: Mean ± SEM. ns p> 0.05, * p<0.05, ** p<0.01 by the Mann-Whitney U-test (panels b, c), or linear mixed-effects modeling with post-hoc comparison (panel d). Multiple comparisons corrected by quantifying the False Discovery Rate and adjusting p-values.

In inflammatory pain, cytokines like TNF-α drive pro-nociceptive ion channels to both somatic and axonal membranes^36,57,61^. To investigate if UbiquiNa_V_ could normalize the delivery and distribution of Na_V_1.8 throughout neuronal compartments in the setting of inflammation, we evaluated the surface expression of Halo-Na_V_1.8 channels after incubation with TNF-α (**Figure 7a**). As observed previously, incubation of neurons with TNF-α resulted in an accumulation of Na_V_1.8 channels at both somatic and axonal membranes. UbiquiNa_V_-expressing neurons exposed to TNF-α had greatly reduced expression of channels in both the cell body (838.9 ± 132.5 vs. 3787.0 ± 935.2 A.U., p<0.05; **Figure 7b**) and at distal axons (74.4 ± 8.3 vs. 293.8 ± 76.9 A.U., p<0.01; **Figure 7c**).

**Figure 7.**
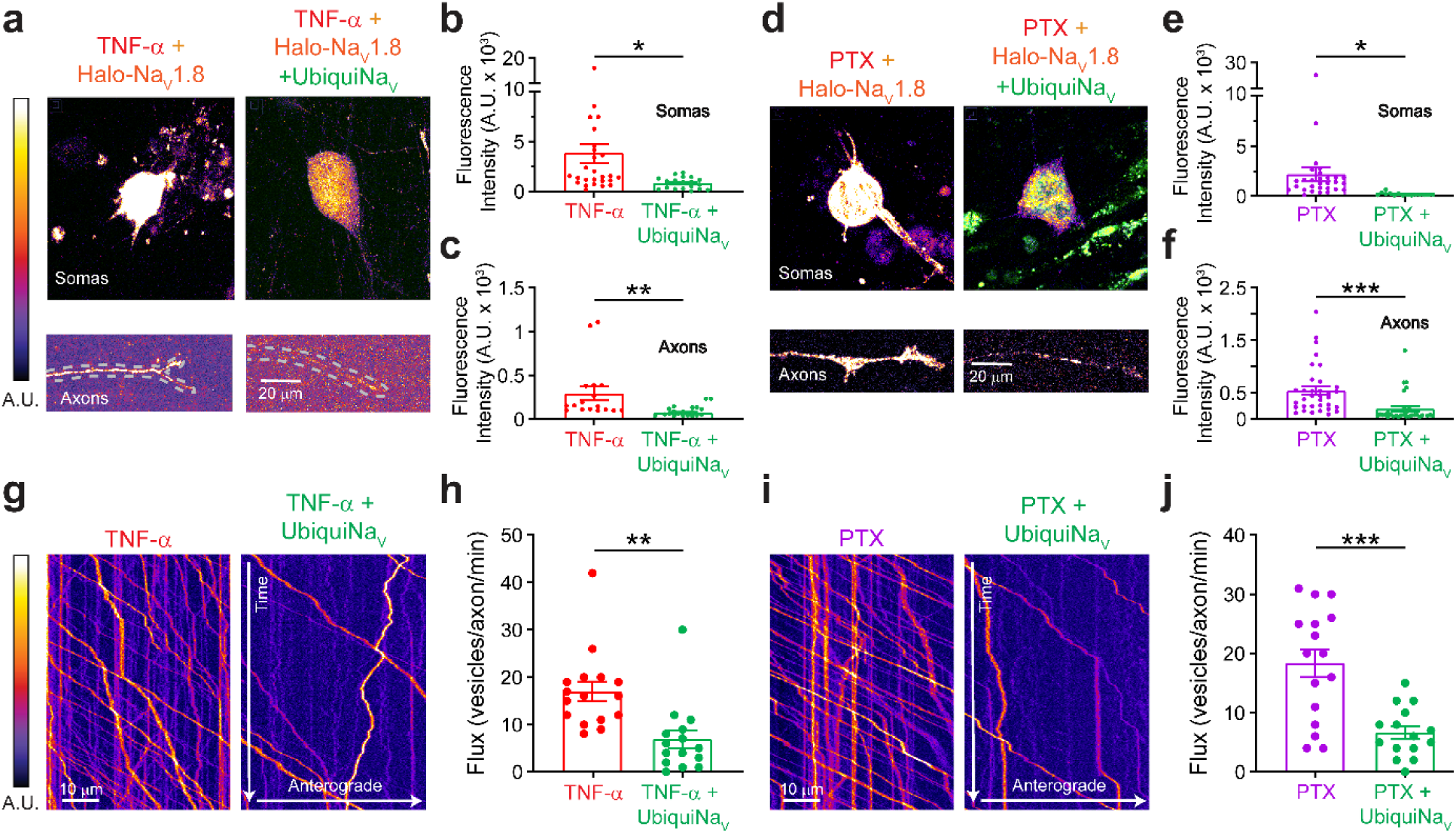
UbiquiNa_V_ normalizes the delivery and distribution of Na_V_1.8 channels in *in vitro* models of inflammatory and chemotherapy-induced neuropathic pain. (a) Representative images of Halo-Na_V_1.8 expression at neuronal surfaces in Halo-Na_V_1.8-expressing neurons incubated with 40 ng/mL TNF-α transfected with eGFP-control (left panels) or UbiquiNa_V_ (right panels). Top panels – somatic expression. Bottom panels – Axonal expression. Channels at the soma and axonal plasma membrane were detected by the HaloTag ligand JF635i. (b) Quantification of somatic Halo-Na_V_1.8 surface expression in TNF-α incubated neurons transfected with Halo-Na_V_1.8-P2A-mRuby2 and eGFP-control (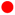; n=27) or UbiquiNa_V_ (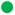; n=19). (c) Quantification of axonal Halo-Na_V_1.8 surface expression in TNF-α incubated neurons transfected with Halo-Na_V_1.8-P2A-mRuby2 and eGFP-control (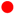; n=17) or UbiquiNa_V_ (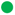; n=20). (d) Representative images of Halo-Na_V_1.8 expression at neuronal surfaces in Halo-Na_V_1.8-expressing neurons incubated with 50 nM Paclitaxel transfected with eGFP-control (left panels) or UbiquiNa_V_ (right panels). Top panels – somatic expression. Bottom panels – Axonal expression. (e) Quantification of somatic Halo-Na_V_1.8 surface expression in TNF-α incubated neurons transfected with Halo-Na_V_1.8-P2A-mRuby2 and eGFP-control (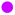; n=29) or UbiquiNa_V_ (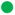; n=15). (f) Quantification of axonal Halo-Na_V_1.8 surface expression in TNF-α incubated neurons transfected with Halo-Na_V_1.8-P2A-mRuby2 and eGFP-control (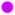; n=34) or UbiquiNa_V_ (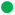; n=30). (g) Representative Kymographs from an axon containing anterogradely trafficking vesicles carrying Halo-Na_V_1.8 in neurons incubated with TNF-α and transfected with eGFP-control or UbiquiNa_V_. (h) Quantitation of anterograde vesicular flux of Halo-Na_V_1.8 channels in neurons incubated with TNF-α and transfected with eGFP-control (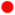; n=16) or UbiquiNa_V_ (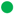; n=15). (i) Representative Kymographs from an axon containing anterogradely trafficking vesicles carrying Halo-Na_V_1.8 in neurons incubated with Paclitaxel and transfected with eGFP-control or UbiquiNa_V_. (j) Quantitation of anterograde vesicular flux of Halo-Na_V_1.8 channels in neurons incubated with Paclitaxel and transfected with eGFP-control (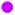; n=16) or UbiquiNa_V_ (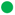; n=15). Bars: Mean ± SEM. ns p> 0.05, * p<0.05, ** p<0.01, *** p<0.001 by the Mann-Whitney U-test (panels b, c, e, f, h), or Student’s unpaired t-test (panel j).

Microtubule inhibiting chemotherapeutic agents like Paclitaxel (PTX) frequently lead to the development of neuropathic pain^62,63^. One of the mechanisms by which neuropathic pain may develop in response to these agents is increased delivery of pro-nociceptive ion channels like Na_V_1.7 and Na_V_1.8 to distal axons^35,64^. To evaluate whether UbiquiNa_V_ expression is protective against Na_V_1.8 accumulation in response to PTX treatment, we evaluated the surface expression of Halo-Na_V_1.8 channels following 48 hours of incubation with 50 nM PTX in neurons expressing UbiquiNa_V_ vs. eGFP-control (**Figure 7d**). UbiquiNa_V_ expressing neurons demonstrated markedly lower surface expression of Halo-Na_V_1.8 at both somas (194.4 ± 39.44 vs. 2187.0 ± 689.3 A.U., p<0.05; **Figure 7e**) and axons (200.3 ± 52.2 vs 545.0 ± 78.8 A.U., p<0.001; **Figure 7f**) in response to PTX treatment.

We also studied trafficking of Na_V_1.8 in these hyperexcitable states and found that TNF-α incubation increases vesicular delivery of Na_V_1.8 to distal axons (**Figure 7g**). UbiquiNa_V_ expression significantly reduces the flux of vesicles carrying Halo-Na_V_1.8 to the distal axon following TNF-α incubation (6.9 ± 1.9 vs. 17.0 ± 2.1 vesicles/axon/min, p<0.01; **Figure 7h**). We observed similar results in neurons incubated with PTX (**Figure 7i**) - vesicular trafficking of Halo-Na_V_1.8 to distal axons was significantly reduced in neurons expressing UbiquiNa_V_ post-PTX treatment (18.4 ± 2.4 vs. 6.7 ± 1.1 vesicles/axon/min, p<0.001; **Figure 7j**).

## Discussion

In this study, we describe a novel strategy for reducing excitability, and a novel agent, UbiquiNa_V_, that functions as a selective regulator of Na_V_1.8 channels in sensory neurons of the dorsal root ganglia. UbiquiNa_V_ leverages the isoform selectivity of a known intracellular binder of Na_V_1.8 (**Figure 1**) and links it to the catalytic activity of a classical E3 ubiquitin ligase (**Figure 2**). This powerful combination enables the post-translational reduction of Na_V_1.8 channels at the cell surface of DRG neurons. We show that UbiquiNa_V_ reduces Na_V_1.8 current density by reducing surface expression of the channel (**Figure 3**). Importantly, we demonstrate that UbiquiNa_V_ is selective for Na_V_1.8 over all mammalian Na_V_ channels (**Figure 4**), and show using two parallel approaches that UbiquiNa_V_ does not have off-target effects on any components of the neuronal electrogenisome in sensory neurons (**Figure 5**). UbiquiNa_V_ acts to decrease Na_V_1.8 surface expression at both the neuronal cell body as well as the distal axon and UbiquiNa_V_ proteins fail to travel to distal axons in the absence of Na_V_1.8, further indicating associative selectivity (**Figure 3**; **Supplementary** Figure 1). The main action of UbiquiNa_V_ appears to take place in the soma, as vesicular delivery of Na_V_1.8 channels to the distal axon is greatly dampened in the presence of UbiquiNa_V_. Through this activity, UbiquiNa_V_ can normalize distribution and delivery of Na_V_1.8 channels throughout the complex trajectory of the sensory neuron and its axon. Our results demonstrate that this strategy normalizes neuronal hyperexcitability (**Figure 6**) under conditions that recapitulate aspects of pain induced by inflammation and chemotherapy (**Figure 7**), an important consideration for *in vivo* analgesia.

Small molecule Na_V_1.8 blockers have reached statistically significant endpoints in clinical trials for acute pain^16^, though benefits in chronic pain syndromes have not been reported to date. The delivery of these molecules (need to cross the blood-nerve barrier, continuous oral redosing, etc.) has been challenging, and this may contribute to relatively small clinical effect sizes of these agents^15,19^. Novel approaches, such as gene therapy, are emerging as potential analgesic agents due to significant advantages over small molecule blockers, such as targeted tissue delivery and extended duration of action^65–68^. Our results demonstrate that a therapeutic heterobifunctional protein, UbiquiNa_V_, achieves selective reduction in the cellular distribution of Na_V_1.8 with concomitant effects on neuronal hyperexcitability under conditions that mimic painful disease states. These results provide strong support for the development of this approach as an analgesic therapy. Importantly, our strategy provides a scaffold for the development of other heterobifunctional reagents aimed at other excitability disorders, for example epilepsy, depending on the discovery of selective binders of the target channel.

We acknowledge several limitations of this study. First, though we investigated modulation of excitability in rodent neurons after treatment with inflammatory and chemotherapy agents, significant species differences exist in both Na_V_ channel expression as well as biophysics^56^, thus studies in native human DRG neurons are needed to further strengthen the translational value of these investigations. Additionally, UbiquiNa_V_ expression did not totally eliminate Na_V_1.8 currents. This may, however, provide an advantage when compared to a CRISPR/Cas9 approach to gene silencing, since some Na_V_1.8 channels will remain, so that nociceptive firing (and the protection it confers) may still be possible. Indeed, our results demonstrate that the main action of UbiquiNa_V_ is to normalize hyperexcitability, not ablate neuronal firing entirely. Our *in vitro* results suggest that delivery of UbiquiNa_V_ DNA to sensory neurons should provide analgesia in *in vivo* pain models. A critical element to this putative therapeutic strategy is transduction of a sufficient number of neurons *in vivo* to permit a therapeutic effect. Future studies will aim to effectively deliver anti-Na_V_ bioPROTACs like UbiquiNa_V_ to sensory neurons *in vivo* to evaluate true analgesic effect.

To our knowledge, we are the first to demonstrate functional reduction in nociceptor activity in response to targeted ubiquitination of a Na_V_ channel. Our results demonstrate that it is possible to achieve exquisite selectivity in targeting a Na_V_ channel, and efficacy in suppressing neuronal excitability under conditions that mimic aspects of inflammation and chemotherapy induced neuropathy. Recently, neuronal calcium (Ca_V_) channels were targeted for ubiquitination using functionalized nanobodies^23,24^. Disruption of these Ca_V_ channels resulted in a total abrogation of neuronal calcium current and decreased pain behaviors in mice^65^. These studies suggest that targeted channel degradation may be a realistic approach to the treatment of pain. In contrast to the nociceptive Na_V_ isoforms, which are preferentially expressed in peripheral sensory neurons, the calcium channels targeted in these prior studies are responsible for synaptic vesicle release in a variety of neuronal types^69^. Thus, targeting Na_V_ channels like Na_V_1.8 may provide a therapeutic opportunity that is more specific for the pain pathway.

Na_V_1.8 channels carry the bulk of the current underlying the action potential upstroke in nociceptive neurons^12,14^. Though Na_V_1.7 channels are responsible for subthreshold amplification, Na_V_1.8 channel open probability at the average AP threshold is ∼500-fold higher than Na_V_1.7^70^. Even in situations where gain-of-function of Na_V_1.7 drives neuronal hyperexcitability, Na_V_1.8 represents 90% of the total sodium influx at the AP threshold, and reduction of Na_V_1.8 can normalize hyperexcitability. A 25-50% reduction in Na_V_1.8 conductance (less than was achieved with UbiquiNa_V_ in this study) was sufficient to reverse the hyperexcitability of DRG neurons expressing a gain-of-function Na_V_1.7 mutation that causes pain in humans^70^. Such results suggest that Na_V_1.8 channels are an indication-agnostic target for pain-relief. However, clinical trials of small molecule Na_V_1.8 blockers have shown differential responses in varying pain conditions. Suzetrigine demonstrated a 1-1.5 point reduction in acute post-surgical pain^16^; however, in in lumbosacral radiculopathy, a chronic pain condition, Suzetrigine demonstrated no effect above placebo^20^. Suzetrigine is a state-dependent inhibitor of Na_V_1.8. Strikingly, Suzetrigine, along with other Na_V_1.8 small molecule blockers^71–73^, demonstrate “reverse use dependence”, where depolarization dramatically relieves channel inhibition. This mechanism of action could be unfavorable since in chronic pain conditions, where repetitive nociceptor firing might result in a channel population where small molecules can no longer bind. Additionally, post-translational modifications of Na_V_ channels in chronic pain disorders may play a role in differentiating acute vs. chronic pathologies^28,74,75^. The biophysical properties of Na_V_1.8^70^ coupled with its favorable expression profile position the channel as a very high-value and clinically validated^16^ target for the treatment of pain; however, there may be a need for alternative mechanisms to small molecule inhibitors for unlocking its full therapeutic potential, particularly in chronic contexts.

Degron-tagged Na_V_ channel constructs were shown to be amenable to degradation by commercially available small-molecules that recruit the E3 ubiquitin ligases Von Hippel Lindau and Cereblon^76^. These results demonstrate that expansion of the therapeutic Na_V_-targeting space to technologies that utilize targeted protein ubiquitination is possible. UbiquiNa_V_ serves as a prototype for isoform-specific membrane protein targeting, such as ion channels. Designing small-molecule PROTACs for Na_V_ channels is challenging due to their high sequence and structural similarity and known small-molecule binders failed as E3 ligase warheads^76^. However, peptides that bind with isoform specificity are easier to identify, and we demonstrate that their binding modules can be leveraged for protein ubiquitination. In the future, these proteins may even be synthesized *de novo*^77^, allowing for customization of target selection and efficacy.

By leveraging the endogenous ability of sensory neurons to regulate the composition of their electrical membranes, we provide a strategy that significantly reduces expression of Na_V_1.8 channels in the plasma membrane and reduces Na_V_1.8 currents in rodent sensory neurons. This strategy selectively targets Na_V_1.8 over other Na_V_ isoforms, normalizes the distribution of Na_V_1.8 protein to distal axons, and normalizes the neuronal hyperexcitability in in vitro models of inflammatory and chemotherapy-induced neuropathic pain. Our results serve as a blueprint for the design of therapeutics that leverage the selective ubiquitination of Na_V_1.8 channels for analgesia.

## Acknowledgements

This work was supported by Merit Review Awards B9253-C and BX004899 from the U.S. Dept. of Veterans Affairs Rehabilitation Research and Development Service and Biomedical Laboratory Research and Development Service, respectively (S.G.W. and S.D.D-H.). The Center for Neuroscience & Regeneration Research is a Collaboration of the Paralyzed Veterans of America with Yale University. S.T. was supported by the NIH/NINDS 1F31NS135909-01. S.T. and G.P.H-R. were supported by NIH/NIGMS Medical Scientist Training Program T32GM007205. M-R.G. is a Banting Fellow and is supported by the Canadian Institutes of Health Research (CIHR). G.P.H-R. was supported by NIH/NINDS 1F31NS122417-01. The content is solely the responsibility of the authors and does not necessarily represent the official views of the National Institutes of Health.

## Author contributions

Conceptualization – S.T., S.D.D-H. Methodology – S.T., M.A, G.P.H-R. Formal Analysis – S.T., M-R.G., M.A., G.P.H-R. Investigation – S.T., M-R.G., M.A., P.E. Resources, F.D-H., P.Z., S.L. Writing, original draft preparation – S.T. Writing, review and editing – S.T., M-R.G., M.A., P.E., G.P.H-R., S.G.W., S.D.D-H. Visualization – S.T. Supervision – S.G.W., S.D.D-H. Project administration – S.T., S.G.W., S.D.D-H. Funding acquisition – S.T., S.G.W., S.D.D-H.

## Methods

### Electrophysiology

#### Rodent Neuron Isolation

Animals used in this study were 2-4 d old Sprague-Dawley rat pups or 4–6-week-old Na_V_1.8-null mice. DRGs or SCGs from these animals were harvested and dissociated as described previously^78–80^. In every culture, neurons were harvested from one female and one male animal so that the number of neurons from each sex were randomly distributed.

#### Rodent DRG Neuron Isolation

Dissected DRGs were incubated at 37° C for 20 min in complete saline solution (CSS) [in mM: 137 NaCl, 5.3 KCl, 1 MgCl2, 25 sorbitol, 3 CaCl2, and 10 HEPES (pH 7.2), adjusted with NaOH], supplemented with 0.6 EDTA and collagenase A (1.5 mg/ml). DRG tissue was then incubated for 20 min at 37° C in CSS containing collagenase D (1.5 mg/mL;), 0.6 EDTA, and papain (30 U/ml). DRG tissue was centrifuged and triturated in 1 ml of DRG culture medium DMEM/F12 (1:1) with penicillin (100 U/ml), streptomycin (0.1 mg/ml), 2 L-glutamine, and 10% FBS containing BSA (1.5 mg/ml) and trypsin inhibitor (1.5 mg/ml). The suspended neurons were then filtered through a 70 μm nylon mesh cell strainer, and then washed with DRG culture medium.

#### Rodent SCG Neuron Isolation

SCGs were dissociated with a 20-minute incubation at 37 °C in oxygenated complete saline solution (CSS) (in mM: 137 NaCl, 5.3 KCl, 1 MgCl2, 25 sorbitol, 3 CaCl2, and 10 HEPES, adjusted to pH 7.2 with NaOH) containing 1.5 mg/ml Collagenase A (Roche, Indianapolis, IN) and 0.6 mM EDTA, followed by an 20-minute incubation at 37 °C in oxygenated CSS containing 1.5 mg/ml collagenase D (Roche), 0.6 mM EDTA and 30 U/ml papain (Worthington Biochemical, Lakewood, NJ). SCGs were then centrifuged and triturated in 0.5 mL of DRG media [DMEM/F12 (1:1) with 100 U/ml penicillin, 0.1 mg/ml streptomycin (Invitrogen), and 10% fetal bovine serum (Hyclone), and 2 mM L-glutamine (Invitrogen)] supplemented with nerve growth factor (NGF, 50 ng/ml) and glial cell line-derived neurotrophic factor (GDNF, 50 ng/ml

#### Electroporation – rat pup DRG neurons

After the preparation of DRG neuron suspension, the cells were centrifuged for 3 min (100×g) and the supernatant was removed. Cells were gently resuspended at room temperature in 100 μL of a mixed Nucleofector solution (82 μL NucleofectorTM solution + 18 μL Supplement). Then, a 100 μL cell suspension with DNA (1.5 μg for all constructs except for eGFP-control (0.3 ug)) was mixed by pipetting. Afterwards, a 20 μL cell suspension was transferred into a strip provided in the Nucleofector kit (Lonza; cat no. V4XP-3032). The sample was covered with the bottom of the strip, and air bubbles were avoided while pipetting. The strip was closed with the cap. The strip was inserted into its holder in the Nucleofector. Program DN-100 was used for rat DRG neurons. After transfection, 60 μL of pre-warmed BSA/TI solution was immediately added, and 80 μL were seeded per coverslip. DRG medium was added to a final volume of 1 mL per well 30 min after seeding, and the cells were incubated at 37° C in 95% O2 and 5% CO2.

#### Electroporation – Na_V_1.8-null mouse neurons

Na_V_1.8-null mouse neurons were transfected with a Nucleofector IIS and Amaxa Basic Neuron SCN Nucleofector Kit. Briefly, following harvest, the cell suspension was centrifuged (100 × g for 3 min), and the cell pellet was resuspended in 20 µL of Nucleofector solution, mixed with the following amounts of DNA depending on the experiment: 1.5 µg for human-Na_V_1.8, 1.5 ug for UbiquiNa_V_, 1.5 ug for Sclt1, 1.5 ug for SCLT1, 0.1 µg mCherry. After electroporation using Nucleofector IIS and protocol SCN-BNP 6, 100 μL of calcium-free DMEM (37° C) was added, and cells were incubated at 37° C for 5 min to recover. The cell mixture was then diluted with DRG media containing 1.5 mg/mL BSA (low endotoxin) and 1.5 mg/mL trypsin inhibitor (Sigma), seeded 100 µl onto poly-D-lysine/laminin-coated coverslips (BD), and incubated at 37° C in a 95% air/5% CO2 (vol/vol) incubator for 45 min to allow neurons to attach to the coverslips. After 45 min, 0.9 mL of DRG media was added into each well, and the DRG neurons were maintained at 37° C in a 95% air/5% CO2 (vol/vol) incubator before voltage-clamp recordings.

#### Electroporation – rat pup SCG neurons

After trituration, neurons were transfected with either 2.5 µg of hNa_V_1.9 WT and 0.1 µg of mCherry or 2.5 µg of hNa_V_1.9 WT and 0.1 µg of mCherry and 1.0 µg of MTD-HECT-p2A GFP using a Nucleofector IIS (Lonza, Basel, Switzerland) and Amaxa Basic Neuron SCN Nucleofector Kit (VSPI—1003) with SCN Basic Neuro Program 6. Transfected neurons were maintained at 37°C in a 95% air/5% CO2 (vol/vol) incubator for 24-48 hrs before voltage-clamp recording (2).

### General Voltage-Clamp

Patch pipettes were fabricated from borosilicate glass (World Precision Instruments) using a P-97 puller (Sutter Instruments) and fire-polished for a resistance of 0.8-1.2 megaohms when filled with internal solution. Unless otherwise noted, the pipette internal solution contained (in mM): 140 CsF, 10 NaCl, 1.1 EGTA, 10 HEPES, 20 Dextrose (pH 7.3 with CsOH, adjusted to 310 mOsm/L with dextrose). External bath solution contained (in mM): 90 NaCl, 50 Choline-Cl, 20 TEA-Cl, 3 KCl, 1 CaCl_2_, 1 MgCl_2_, 10 HEPES, 5 Sucrose, 0.1 CdCl_2_, ± 0.001 TTX (pH 7.3 with NaOH, adjusted to 320 mOsm/L with sucrose).

Macroscopic currents were recorded in voltage-clamp mode using an EPC-10 amplifier and the PatchMaster Next program (HEKA Electronik). Sodium currents were recorded in the whole-cell configuration. Cells with a leak current >200 pA were excluded. Series resistance compensation of 80-90% was applied to reduce voltage error. Cells were excluded if the voltage error exceeded 5 mV. Recordings were sampled at 50 kHz through a low-pass Bessel filter of 10 kHz. After achieving the whole-cell configuration, a 5-min delay was applied to allow adequate time for the pipette solution and cytoplasmic milieu to equilibrate. Unless otherwise noted, Cells were held at −80 mV.

I-V relationships generated by the voltage protocol were fit according to the following equation:

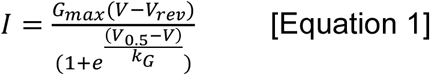

#### Na_V_1.8 recordings from DRG neurons

Transfected DRG neurons with diameters between 20-30 uM were selected for recording. Positively transfected neurons were identified by green (for eGFP linked constructs) or red (for mRuby or mCherry linked constructs) fluorescence. After achieving the whole-cell configuration, neurons were held at −80 mV. To isolate TTX-R Na_V_1.8 currents, we included 1 uM TTX in the bath. To evoke currents through Na_V_1.8, we used a standard activation protocol. Briefly, we applied a series of 100ms step depolarizations from −80 to +40 mV in 5-mV increments applied at 10s interpulse intervals.

### TTX-S Current Recordings from DRG neurons

Transfected DRG neurons between 20-30 μm were selected for recording. The internal pipette solution was the standard voltage clamp solution, but the external solution did not contain TTX. After achieving the whole-cell configuration, neurons were held at −90 mV (a potential at which most TTX-S channels are not inactivated). Then, to record total sodium currents, a series of 100ms step depolarizations from −100 to +40 mV in 5-mV increments were applied at 10s interpulse intervals. After this set of data was recorded, the holding potential was switched to −50 mV (a potential at which predominantly Na_V_1.8 channels are active) and the same voltage protocol was applied to the cell. The TTX-S component of the total sodium current was determined by reference subtraction of the recordings at −50 mV from those at −90 mV. IV curves were plotted and fit with Equation 1. Peak current density was defined as previously described.

#### hNa_V_1.9 recordings from SCG neurons

Na_V_1.9 currents were recorded in rat pup SCG neurons, which do not express native TTX-R currents. External bath solution for these recordings contained 140 NaCl, 20 TEA-Cl, 3 KCl, 1 CaCl_2_, 1 MgCl_2_, 10 HEPES, 5 Sucrose, 0.1 CdCl_2_, 0.001 TTX (pH 7.3 with NaOH, adjusted to 320 mOsm/L with sucrose).

### Flow sorting

Suspension HEK293 cells (Expi293F) were sourced from ThermoFisher. ExpiHEK cells were transfected with Gibco Expifectamine transfection kit following manufacturer protocols. Briefly, cells at a density of ∼3 x 10^6^/mL were transfected with Na_V_ channel constructs (tagged with -P2A-eGFP) and either UbiquiNa_V_-P2A-mCherry or mCherry. Transfection enhancing solution was added to cell suspensions 24 hours after plasmid addition as per manufacturer protocol. 48 hours after plasmid addition, cells were harvested for FACS.

For ND7/23 cells, transfection was achieved with the Lipofectamine 2000 reagent following manufacturer protocols. Cells were transfected with eGFP-2A-Na_V_1.8 and either UbiquiNa_V_-2AmCherry or mCherry. 48 hours later, cells were dissociated using Accutase.

FACS was conducted with cells resuspended in ExCell media containing 5 U/mL DNase I and 5 mM MgCl_2_ using a Wolf G2 NanoCellect Cell Sorter. Sheath fluid was identical to sample fluid. A minimum of 500,000 cells were collected in each condition. After FACS, samples were manually inspected to ensure >95% of cells were doubly positive for green and red fluorescence.

#### Automated Electrophysiology – Heterologous Systems

Cells were resuspended in 3 mL of extracellular recording solution (see below) before placement in the 1 well Cell Transfer Plate. Sodium currents were measured in the whole-cell configuration using a Qube-384 (Sophion A/S, Copenhagen, Denmark) automated voltage-clamp system. Intracellular solution contained (in mM): 120 CsF, 10 NaCl, 2 MgCl2, 10 HEPES, adjusted to pH 7.2 with CsOH. The extracellular recording solution for experiments with Expi293 cells contained (in mM): 145 NaCl, 3 KCl, 1 MgCl2, 1.5 CaCl2, 10 HEPES, adjusted to pH 7.4 with NaOH. The extracellular recording solution for experiments with ND7/23 cells contained 70 NaCl, 70 Choline-Cl, 20 TEA-Cl, 3 KCl, 1 CaCl_2_, 1 MgCl_2_, 10 HEPES, 5 Sucrose, 0.1 CdCl_2_, 0.001 TTX (pH 7.3 with NaOH, adjusted to 320 mOsm/L with sucrose). Liquid junction potentials calculated to be ∼7 mV were not adjusted for. Currents were low pass filtered at 5 kHz and recorded at 25-kHz sampling frequency. Series resistance compensation was applied at 100% and leak subtraction enabled. The Qube-384 temperature controller was used to maintain recording chamber temperature for all experiments at 22 ± 2°C at the recording chamber. Appropriate filters for cell membrane resistance (typically >500 MOhm), series resistance (<10 MOhm), and Na_V_ current magnitude (>500 pA at a test pulse from a resting HP of −120 mV) were routinely applied to exclude poor quality recordings.

The experimenter responsible for conducting automated patch clamp experiments and analysis was blinded to the conditions of the transfection. A separate experimenter assigned labels to the data after unblinding.

#### Current clamp electrophysiology

Recordings from rodent DRG neurons were taken 24-48 hours post-transfection. Extracellular bath solution contained (in mM): 140 NaCl, 3 KCl, 2 CaCl2, 2 MgCl2, 15 dextrose, and 10 HEPES. Osmolarity was brought to approximately 320 mOsm with sucrose and the pH was titrated to 7.3 with NaOH. Intracellular pipette solution contained (in mM): 140 KCl, 3 Mg-ATP, 0.5 EGTA, 5 HEPES, and 20 dextrose. Osmolarity was similarly adjusted to approximately 320 mOsm and the pH was adjusted to 7.3 using KOH. Current-clamp recordings were sampled at 50 KHz and filtered using two Bessel filters at 10 and 2.9 KHz.

DRG neurons with an input resistance lower than 100 MΩ were excluded from analysis. Input resistance was determined by the slope of a linear fit to hyperpolarizing responses to current steps from −5 pA to −40 pA in −5 pA increments. Action potential repetitive firing was determined by summing the total number of action potentials that a neuron fired after a 1 s current injection.

Current threshold was defined as the first current injection step that resulted in action potential firing without subsequent failure and was determined by a series of depolarizing current injections (200 ms) that increased incrementally by 5 pA. For the calculation of threshold, action potentials were defined as rapid increases in membrane potential to >40 mV with a total amplitude >80 mV. However, as neurons often attenuate firing with repetitive action potential spiking, when examining repetitive firing, action potentials were counted if the membrane potential rapidly crossed 0 mV, regardless of overshoot or total amplitude. Action potential repetitive firing was determined by summing the total number of action potentials that a neuron fired after a 1 s current injection.

Dynamic clamp recordings of Na_V_1.8-null DRG neurons

Current clamp procedures were the same as described above. DRG neurons from Na_V_1.8-null mice were harvested and transfected as described.

A Hodgkin & Huxley kinetic model of Na_V_1.8 was derived from published literature^56^. DRG neurons were dynamically clamped using the Cybercyte DC1 dynamic clamp system (Cytocybernetics, Buffalo NY). The amount of Na_V_1.8 current injected into each neuron was specific to the capacitance of each cell, as described in previous literature^81^

In brief, the Na_V_1.8 channel model was based on Hodgkin-Huxley equations:

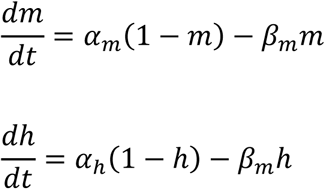

where m and h represent channel activation and inactivation gates, and α and β are forward and backward rate constants, respectively. These rate constants were voltage-dependent and defined by the following equations:

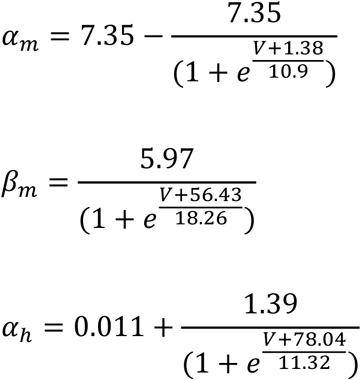

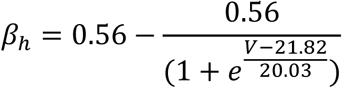

### Imaging

As described previously^34,36,37,57,61^, MFCs were bound to glass-bottomed dishes according to manufacturer’s instructions. Dissected DRGs were transfected as described previously ^37,82^ and plated into the somatic chamber of prepared MFCs containing DRG media with growth factors. Media in the axonal chambers was supplemented with 2x growth factors. After 24 hours, medium was changed to serum free-medium in both chambers. After 5-7 days in vitro, DRG neurons were taken for imaging experiments. For experiments involving TNF-α or Paclitaxel, neurons were incubated in either 20 ng/mL TNF-α, 50 nM PTX, or DMSO control for 24 hours prior to imaging for TNF-α experiments, or 48 hours prior to imaging for PTX experiments.

Neurons were washed with DRG-Neuronal Imaging Saline (NIS) for these experiments. DRG-NIS contained (in mM): 136 NaCl, 3 KCl, 1 MgSO4, 2.5 CaCl2, 0.15 NaH2PO4, 0.1 Ascorbic Acid, 20 HEPES, 8 Dextrose (pH 7.4 with NaOH, adjusted to 320 mOsm/l)

#### Surface Expression

Halo-Na_V_1.8 channels at the somatic and axonal surface were labeled by incubation with cell-impermeable JF635i-HaloTag ligand. Both somatic and axonal compartments were thoroughly washed with DRG-NIS, fixed with 4% paraformaldehyde, and then taken for confocal fluorescence imaging.

We measured the fluorescence signal intensity of surface-labeled Halo-Na_V_1.8 from compressed confocal z-stacks of the distal 50 μM of axons.

#### Trafficking – OPAL imaging

Trafficking assays in these studies were variations on the Optical Pulse-chase Axonal Long-distance (OPAL) imaging method described previously^36,57,61^. 100 nm cell-permeable JFX650-Halo ligand was applied to the somatic chamber of MFCs containing DRG neurons transfected with Halo-Na_V_1.8 and UbiquiNa_V_ constructs. After 15 minutes of incubation, the chambers were thoroughly washed with NIS, and MFCs were placed in a 37° C stage-top incubator for imaging. Halo-Na_V_1.8 positive neurons were identified by red fluorescence (from the mRuby2 reporter) and UbiquiNa_V_ positive neurons were identified by green fluorescence (eGFP reporter). Multiple fields of view in the axonal chamber were selected, and anterogradely trafficking vesicles were imaged in the far-red channel continuously for 2 minutes.

Resulting movies were opened in ImageJ and the KymographClear toolset was used to create kymographs of the selected axons^83^. Axons containing anterogradely-moving vesicles were traced manually using a 50 μm segmented line, and the signal under that line was converted into a two-dimensional image that could be further analyzed. KymoButler is a machine learning algorithm that traces vesicle tracks from kymographs^84^. Vesicle flux was determined by counting the number of vesicles which crossed the midline of the kymograph. Vesicle intensity was determined as the average of fluorescence values of pixels along the vesicle track, minus the background signal of the kymograph (defined as the modal value for that kymograph). The fluorescence intensities of multiple vesicles within axons were averaged.

#### Cotrafficking – OPAL imaging

Cotrafficking assays in these studies were variations on the Optical Pulse-chase Axonal Long-distance (OPAL) imaging method described previously^34,36,37^. 100 nm cell-permeable JFX650-Halo ligand and 100 nm cell-permeable JFX554cp-SNAP ligand were applied to the somatic chamber of MFCs containing DRG neurons from Na_V_1.8-null mice transfected with Halo-Na_V_1.8 and SNAP-UbiquiNa_V_-C942S constructs.

Anterogradely moving vesicles containing SNAP-UbiquiNa_V_-C942S were identified, and axons within the axonal chamber were then imaged in red and far-red by rapid laser and color filter switching. Kymographs were generated as above, and kymographs from each fluorescent channel were analyzed to score vesicles as positive for one or both proteins. The background was measured for each color and kymograph and subtracted from the vesicle measurements. A cutoff of 100 A.U. was used to categorize vesicles as positive or negative for each protein.

### Molecular Biology

#### Plasmids

Sclt1: Rat Sclt1 (also known as CAP-1A) was previously reported^27^. Briefly, the complete coding sequence (688 a.a.) was subcloned in-frame into the NheI/BamHI sites of pEGFP-N1 vector (Clontech) to produce plasmid pCAP-1A-eGFP. Full-length Rat SCLT1 cDNA generated by double digestion of pCAP-1A-eGFP with NheI/AgeI was subcloned into the XbaI/AgeI sites of pcDNA3.1B (Invitrogen), and the native translation termination codon was restored by site directed mutagenesis using the QuickChange system (Stratagene, La Jolla, CA) to produce the vector pcDNA3-CAP-1A (Referred to as Sclt1 hereinafter).

Human Na_V_1.8: The codon-optimized human Na_V_1.8 construct (pcDNA5-SCN10A) was purchased from Genionics, and previously reported^79^. Codon-optimized Halo-Na_V_1.8 construct was previously reported^34^. The final construct topology is in order from the N-terminus:1-30 a.a. β 4 signal peptide, 3 myc tag (EQKLISEEDL), Halo-tag enzyme (297 a.a.) (Promega), 3 HA tag (YPYDVPDYA), 30 a.a. extracellular stalk (β 4 132-162), 21 a.a. transmembrane segment (β 4 163-183), 7 a.a. linker (SGLRSAT), hNa_V_1.8.

### Statistics

Unless noted otherwise, we performed the following two-stage statistical procedure for all group comparisons. First, we performed testing for normality using the Shapiro-Wilks test and the Kolmogorov-Smirnov test. If the underlying data distribution was normal, we performed two-tailed parametric statistical testing using Student’s t-test. If the distribution of data was non-normal, we compared groups using two-tailed non-parametric testing using the Mann-Whitney U-test^85^. For repeated measures data, we compared groups using mixed effects modeling. A level of significance α=0.05 was used with p-values less than 0.05 considered to be statistically significant. When multiple comparisons were made, we quantified the False Discovery Rate and adjusted p-values accordingly to correct for multiple comparisons. All values are reported as means ± standard error of means (SEM).

## References

1. Pizzo, P. A. & Clark, N. M. Alleviating Suffering 101 — Pain Relief in the United States. N Engl J Med 366, 197–199 (2012).

2. Johannes, C. B., Le, T. K., Zhou, X., Johnston, J. A. & Dworkin, R. H. The Prevalence of Chronic Pain in United States Adults: Results of an Internet-Based Survey. The Journal of Pain 11, 1230–1239 (2010).

3. Breivik, H., Collett, B., Ventafridda, V., Cohen, R. & Gallacher, D. Survey of chronic pain in Europe: Prevalence, impact on daily life, and treatment. European Journal of Pain 10, 287–287 (2006).

4. Institute of Medicine (US) Committee on Advancing Pain Research, Care, andEducation. Relieving Pain in America: A Blueprint for Transforming Prevention, Care, Education, and Research. (National Academies Press (US), Washington (DC), 2011).

5. Alsaloum, M., Higerd, G. P., Effraim, P. R. & Waxman, S. G. Status of peripheral sodium channel blockers for non-addictive pain treatment. Nat Rev Neurol 16, 689– 705 (2020).

6. Alsaloum, M. et al. Voltage-gated sodium channels in excitable cells as drug targets. Nat Rev Drug Discov 1–21 (2025) doi:10.1038/s41573-024-01108-x.

7. Bennett, D. L., Clark, A. J., Huang, J., Waxman, S. G. & Dib-Hajj, S. D. The Role of Voltage-Gated Sodium Channels in Pain Signaling. Physiological Reviews 99, 1079–1151 (2019).

8. Rush, A. M., Cummins, T. R. & Waxman, S. G. Multiple sodium channels and their roles in electrogenesis within dorsal root ganglion neurons. The Journal of Physiology 579, 1–14 (2007).

9. Akopian, A. N., Sivilotti, L. & Wood, J. N. A tetrodotoxin-resistant voltage-gated sodium channel expressed by sensory neurons. Nature 379, 257–262 (1996).

10. Akopian, A. N. et al. The tetrodotoxin-resistant sodium channel SNS has a specialized function in pain pathways. Nat Neurosci 2, 541–548 (1999).

11. Choi, J.-S. & Waxman, S. G. Physiological interactions between Nav1.7 and Nav1.8 sodium channels: a computer simulation study. Journal of Neurophysiology 106, 3173–3184 (2011).

12. Blair, N. T. & Bean, B. P. Roles of tetrodotoxin (TTX)-sensitive Na+ current, TTX-resistant Na+ current, and Ca2+ current in the action potentials of nociceptive sensory neurons. J Neurosci 22, 10277–10290 (2002).

13. Blair, N. T. & Bean, B. P. Role of Tetrodotoxin-Resistant Na+ Current Slow Inactivation in Adaptation of Action Potential Firing in Small-Diameter Dorsal Root Ganglion Neurons. J Neurosci 23, 10338–10350 (2003).

14. Renganathan, M., Cummins, T. R. & Waxman, S. G. Contribution of Nav1.8 Sodium Channels to Action Potential Electrogenesis in DRG Neurons. Journal of Neurophysiology 86, 629–640 (2001).

15. Waxman Stephen G. Targeting a Peripheral Sodium Channel to Treat Pain. New England Journal of Medicine 389, 466–469 (2023).

16. Jones, J. et al. Selective Inhibition of NaV1.8 with VX-548 for Acute Pain. New England Journal of Medicine 389, 393–405 (2023).

17. Vertex Announces Positive Results From the VX-548 Phase 3 Program for the Treatment of Moderate-to-Severe Acute Pain | Vertex Pharmaceuticals. https://investors.vrtx.com/news-releases/news-release-details/vertex-announces-positive-results-vx-548phase-3-program.

18. Commissioner, O. of the. FDA Approves Novel Non-Opioid Treatment for Moderate to Severe Acute Pain. FDA https://www.fda.gov/news-events/press-announcements/fda-approves-novel-non-opioid-treatment-moderate-severe-acute-pain (2025).

19. Wallace, M. S. Trials for Managing Acute Pain — A Clinically Meaningful Small Effect Size? New England Journal of Medicine 389, 464–465 (2023).

20. Vertex Announces Results From Phase 2 Study of Suzetrigine for the Treatment of Painful Lumbosacral Radiculopathy | Vertex Pharmaceuticals. https://investors.vrtx.com/news-releases/news-release-details/vertex-announces-results-phase-2-study-suzetrigine-treatment.

21. Békés, M., Langley, D. R. & Crews, C. M. PROTAC targeted protein degraders: the past is prologue. Nat Rev Drug Discov 21, 181–200 (2022).

22. Kanner, S. A., Morgenstern, T. & Colecraft, H. M. Sculpting ion channel functional expression with engineered ubiquitin ligases. eLife 6, e29744 (2017).

23. Morgenstern, T. J., Park, J., Fan, Q. R. & Colecraft, H. M. A potent voltage-gated calcium channel inhibitor engineered from a nanobody targeted to auxiliary CaVβ subunits. eLife 8, e49253 (2019).

24. Morgenstern, T. J. et al. Selective posttranslational inhibition of CaVβ1-associated voltage-dependent calcium channels with a functionalized nanobody. Nat Commun 13, 7556 (2022).

25. Marei, H. et al. Antibody targeting of E3 ubiquitin ligases for receptor degradation. Nature 610, 182–189 (2022).

26. VanDyke, D., Taylor, J. D., Kaeo, K. J., Hunt, J. & Spangler, J. B. Biologics-based degraders — an expanding toolkit for targeted-protein degradation. Curr Opin Biotechnol 78, 102807 (2022).

27. Liu, C. et al. CAP-1A is a novel linker that binds clathrin and the voltage-gated sodium channel Nav1.8. Molecular and Cellular Neuroscience 28, 636–649 (2005).

28. Laedermann, C. J. et al. Dysregulation of voltage-gated sodium channels by ubiquitin ligase NEDD4-2 in neuropathic pain. J Clin Invest 123, 3002–3013 (2013).

29. Gasser, A. et al. Two Nedd4-binding motifs underlie modulation of sodium channel Nav1.6 by p38 MAPK. J Biol Chem 285, 26149–26161 (2010).

30. Edwin, F., Anderson, K. & Patel, T. B. HECT Domain-containing E3 Ubiquitin Ligase Nedd4 Interacts with and Ubiquitinates Sprouty2. J Biol Chem 285, 255–264 (2010).

31. Foot, N., Henshall, T. & Kumar, S. Ubiquitination and the Regulation of Membrane Proteins. Physiol Rev 97, 253–281 (2017).

32. Raikwar, N. S., Vandewalle, A. & Thomas, C. P. Nedd4–2 interacts with occludin to inhibit tight junction formation and enhance paracellular conductance in collecting duct epithelia. Am J Physiol Renal Physiol 299, F436–F444 (2010).

33. Dubin, A. E. & Patapoutian, A. Nociceptors: the sensors of the pain pathway. J Clin Invest 120, 3760–3772 (2010).

34. Higerd-Rusli, G. P. et al. Depolarizing NaV and Hyperpolarizing KV Channels Are Co-Trafficked in Sensory Neurons. J. Neurosci. 42, 4794–4811 (2022).

35. Baker, C. A. et al. Paclitaxel effects on axonal localization and vesicular trafficking of NaV1.8. Frontiers in Molecular Neuroscience 16, (2023).

36. Akin, E. J. et al. Building sensory axons: Delivery and distribution of NaV1.7 channels and effects of inflammatory mediators. Sci Adv 5, eaax4755 (2019).

37. Tyagi, S. et al. Conserved but not critical: Trafficking and function of NaV1.7 are independent of highly conserved polybasic motifs. Frontiers in Molecular Neuroscience 16, (2023).

38. Goodwin, G. & McMahon, S. B. The physiological function of different voltage-gated sodium channels in pain. Nat Rev Neurosci 22, 263–274 (2021).

39. Nassar, M. A. et al. Nociceptor-specific gene deletion reveals a major role for Nav1.7 (PN1) in acute and inflammatory pain. Proc Natl Acad Sci U S A 101, 12706–12711 (2004).

40. Catterall, W. A., Kalume, F. & Oakley, J. C. NaV1.1 channels and epilepsy. J Physiol 588, 1849–1859 (2010).

41. Yu, F. H. & Catterall, W. A. Overview of the voltage-gated sodium channel family. Genome Biology 4, 207 (2003).

42. Li, Z. et al. Structure of human Nav1.5 reveals the fast inactivation-related segments as a mutational hotspot for the long QT syndrome. Proceedings of the National Academy of Sciences 118, e2100069118 (2021).

43. Wang, J., Ou, S.-W. & Wang, Y.-J. Distribution and function of voltage-gated sodium channels in the nervous system. Channels (Austin) 11, 534–554 (2017).

44. Osorio, N., Korogod, S. & Delmas, P. Specialized functions of Nav1.5 and Nav1.9 channels in electrogenesis of myenteric neurons in intact mouse ganglia. J Neurosci 34, 5233–5244 (2014).

45. Cummins, T. R. et al. A novel persistent tetrodotoxin-resistant sodium current in SNS-null and wild-type small primary sensory neurons. J Neurosci 19, RC43 (1999).

46. Cummins, T. R., Rush, A. M., Estacion, M., Dib-Hajj, S. D. & Waxman, S. G. Voltage-clamp and current-clamp recordings from mammalian DRG neurons. Nat Protoc 4, 1103–1112 (2009).

47. Ghovanloo, M.-R., Tyagi, S., Zhao, P. & Waxman, S. G. Nav1.8, an analgesic target for nonpsychotomimetic phytocannabinoids. Proceedings of the National Academy of Sciences 122, e2416886122 (2025).

48. Sizova, D. V. et al. A 49-residue sequence motif in the C terminus of Nav1.9 regulates trafficking of the channel to the plasma membrane. Journal of Biological Chemistry 295, 1077–1090 (2020).

49. Han, C. et al. Familial gain-of-function Nav1.9 mutation in a painful channelopathy. J Neurol Neurosurg Psychiatry 88, 233–240 (2017).

50. Rush, A. M. et al. A single sodium channel mutation produces hyper- or hypoexcitability in different types of neurons. Proceedings of the National Academy of Sciences 103, 8245–8250 (2006).

51. Waxman, S. G. Sodium channels, the electrogenisome and the electrogenistat: lessons and questions from the clinic. The Journal of Physiology 590, 2601–2612 (2012).

52. Bourinet, E. et al. Calcium-Permeable Ion Channels in Pain Signaling. Physiological Reviews 94, 81–140 (2014).

53. Tsantoulas, C. & McMahon, S. B. Opening paths to novel analgesics: the role of potassium channels in chronic pain. Trends in Neurosciences 37, 146–158 (2014).

54. Vasylyev, D. V., Han, C., Zhao, P., Dib-Hajj, S. & Waxman, S. G. Dynamic-clamp analysis of wild-type human Nav1.7 and erythromelalgia mutant channel L858H. Journal of Neurophysiology 111, 1429–1443 (2014).

55. Alsaloum, M. et al. Contributions of NaV1.8 and NaV1.9 to excitability in human induced pluripotent stem-cell derived somatosensory neurons. Sci Rep 11, 24283 (2021).

56. Han, C. et al. Human Nav1.8: enhanced persistent and ramp currents contribute to distinct firing properties of human DRG neurons. J Neurophysiol 113, 3172–3185 (2015).

57. Tyagi, S. et al. Compartment-specific regulation of NaV1.7 in sensory neurons after acute exposure to TNF-α. Cell Reports 43, 113685 (2024).

58. Ozaktay, A. C. et al. Effects of interleukin-1 beta, interleukin-6, and tumor necrosis factor on sensitivity of dorsal root ganglion and peripheral receptive fields in rats. Eur Spine J 15, 1529–1537 (2006).

59. Chen, X. et al. TNF-α enhances the currents of voltage gated sodium channels in uninjured dorsal root ganglion neurons following motor nerve injury. Exp Neurol 227, 279–286 (2011).

60. Spicarova, D., Nerandzic, V. & Palecek, J. Modulation of spinal cord synaptic activity by tumor necrosis factor α in a model of peripheral neuropathy. J Neuroinflammation 8, 177 (2011).

61. Higerd-Rusli, G. P. et al. Inflammation differentially controls transport of depolarizing Nav versus hyperpolarizing Kv channels to drive rat nociceptor activity. Proceedings of the National Academy of Sciences 120, e2215417120 (2023).

62. 62. Mo, H., et al. Association of Taxane Type With Patient-Reported Chemotherapy-Induced Peripheral Neuropathy Among Patients With Breast Cancer. *JAMA Network Open* 5, e2239788 (2022).

63. Flatters, S. J. L., Dougherty, P. M. & Colvin, L. A. Clinical and preclinical perspectives on Chemotherapy-Induced Peripheral Neuropathy (CIPN): a narrative review. Br J Anaesth 119, 737–749 (2017).

64. Akin, E. J. et al. Paclitaxel increases axonal localization and vesicular trafficking of Nav1.7. Brain 144, 1727–1737 (2021).

65. Sun, L. et al. Targeted ubiquitination of sensory neuron calcium channels reduces the development of neuropathic pain. Proceedings of the National Academy of Sciences 119, e2118129119 (2022).

66. Ovsepian, S. V. & Waxman, S. G. Gene therapy for chronic pain: emerging opportunities in target-rich peripheral nociceptors. Nat Rev Neurosci 1–14 (2023) doi:10.1038/s41583-022-00673-7.

67. Perez-Sanchez, J. et al. A humanized chemogenetic system inhibits murine pain-related behavior and hyperactivity in human sensory neurons. Sci Transl Med 15, eadh3839 (2023).

68. Moreno, A. M. et al. Long-lasting analgesia via targeted in situ repression of NaV1.7 in mice. Sci Transl Med 13, eaay9056 (2021).

69. Dolphin, A. C., Functions of Presynaptic Voltage-gated Calcium Channels. function 2, zqaa027 (2021).

70. Vasylyev, D. V., Zhao, P., Schulman, B. R. & Waxman, S. G. Interplay of Nav1.8 and Nav1.7 channels drives neuronal hyperexcitability in neuropathic pain. Journal of General Physiology 156, e202413596 (2024).

71. Gilchrist, J. M., Yang, N.-D., Jiang, V. & Moyer, B. D. Pharmacologic Characterization of LTGO-33, a Selective Small Molecule Inhibitor of the Voltage-Gated Sodium Channel NaV1.8 with a Unique Mechanism of Action. Molecular Pharmacology 105, 233–249 (2024).

72. Jo, S., Zhang, H.-X. B. & Bean, B. P. Use-Dependent Relief of Inhibition of Nav1.8 Channels by A-887826. Molecular Pharmacology 103, 221–229 (2023).

73. Browne, L. E., Blaney, F. E., Yusaf, S. P., Clare, J. J. & Wray, D. Structural Determinants of Drugs Acting on the Nav1.8 Channel *. Journal of Biological Chemistry 284, 10523–10536 (2009).

74. Bierhaus, A. et al. Methylglyoxal modification of Nav1.8 facilitates nociceptive neuron firing and causes hyperalgesia in diabetic neuropathy. Nat Med 18, 926–933 (2012).

75. 75. Tyagi, S., et al. Compartment-specific regulation of NaV1.7 in sensory neurons after acute exposure to TNF-α. Cell Reports 43, (2024).

76. Chamessian, A., Payne, M., Gordon, I., Zhou, M. & Gereau, R. Small molecule-mediated targeted protein degradation of voltage-gated sodium channels involved in pain. 2025.01.21.634079 Preprint at 10.1101/2025.01.21.634079 (2025).

77. Vázquez Torres, S., et al. De novo designed proteins neutralize lethal snake venom toxins. Nature 1–7 (2025) doi:10.1038/s41586-024-08393-x.

78. Dib-Hajj, S. D. et al. Transfection of rat or mouse neurons by biolistics or electroporation. Nat Protoc 4, 1118–1127 (2009).

79. Faber, C. G. et al. Gain-of-function Nav1.8 mutations in painful neuropathy. Proc Natl Acad Sci U S A 109, 19444–19449 (2012).

80. Han, C. et al. The Domain II S4-S5 Linker in Nav1.9: A Missense Mutation Enhances Activation, Impairs Fast Inactivation, and Produces Human Painful Neuropathy. Neuromol Med 17, 158–169 (2015).

81. Ghovanloo, M.-R. et al. Sodium currents in naïve mouse dorsal root ganglion neurons: No major differences between sexes. Channels 18, 2289256 (2024).

82. Dib-Hajj, S. D. et al. Transfection of rat or mouse neurons by biolistics or electroporation. Nat Protoc 4, 1118–1127 (2009).

83. Higerd-Rusli, G. P. et al. The fates of internalized NaV1.7 channels in sensory neurons: Retrograde cotransport with other ion channels, axon-specific recycling, and degradation. Journal of Biological Chemistry 299, (2023).

84. Jakobs, M. A., Dimitracopoulos, A. & Franze, K. KymoButler, a deep learning software for automated kymograph analysis. eLife 8, e42288 (2019).

85. Rochon, J., Gondan, M. & Kieser, M. To test or not to test: Preliminary assessment of normality when comparing two independent samples. BMC Medical Research Methodology 12, 81 (2012).

