## Supplemental Figures for "Targeted ubiquitination of Na_V_1.8 reduces sensory neuronal excitability"

**Supplementary material for: Targeted ubiquitination of $Na_V1.8$ reduces sensory neuronal excitability**

* Corresponding Authors

Sidharth Tyagi, PhD

Stephen G. Waxman, MD, PhD

Sulayman D. Dib-Hajj, PhD

Neuroscience and Regeneration Research Center, VAMC, 950 Campbell Avenue, Bldg. 34, West Haven, CT 06516


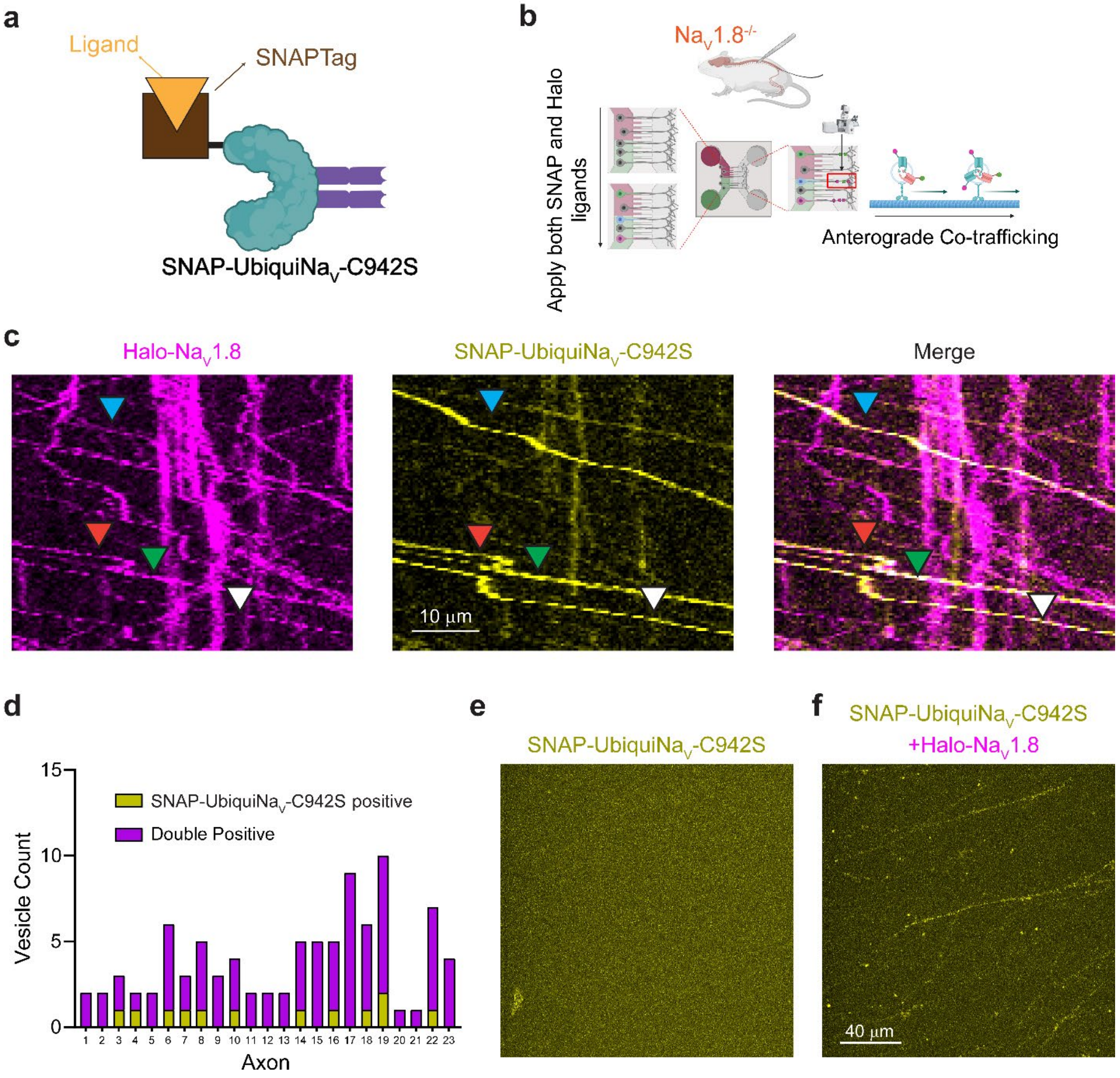


**Supplementary Figure 1. UbiquiNav requires Nav1.8 to travel in vesicles to the distal axon.**

(**a**) SNAP-UbiquiNav-C942S was generated by linking a SNAPTag enzyme to the HECT domain of UbiquiNav. The inactivation of UbiquiNav was necessary since the assay sought to detect UbiquiNav in anterogradely trafficking vesicles. The active UbiquiNav moiety abrogated transport and accumulation of Nav1.8 channels in the soma and axonal compartments and thus was difficult to evaluate by imaging of distal axons.

(**b**) Schematic of Co-trafficking OPAL experiments. DRG neurons from Nav1.8-null mice were transfected with Halo-Nav1.8 and SNAP-UbiquiNav-C942S and plated in MFCs. JFX650-Halo and JFX554cp-SNAP were applied to the somatic chamber of MFCs. After

25 minutes of labeling, MFCs were washed and taken to a confocal microscope for optical pulse-chase imaging of distal axons.

(**c**) Representative kymographs from distal axons of Nav1.8-null neurons carrying Halo-Nav1.8 (left panel) and SNAP-UbiquiNav-C942S (middle panel). Right panel is a merge of the two kymographs. Vesicle tracks positive for SNAP-UbiquiNav-C942S are marked with triangular markers.

(**d**) Co-trafficking of SNAP-UbiquiNav-C942S with Halo-Nav1.8. Axons were analyzed for the presence of vesicles containing SNAP-UbiquiNav-C942S. Vesicles were positive for only SNAP-UbiquiNav-C942S if they did not overlap with fluorescent signal from labeled Halo-Nav1.8 channels.

(**e**) Whole-field image of distal axons of Nav1.8-null neurons expressing SNAP-UbiquiNav-C942S. No JFX554cp-SNAP fluorescence is visible.

(**f**) Whole-field image of distal axons of Nav1.8-null neurons expressing SNAP-UbiquiNav-C942S and Halo-Nav1.8. Fluorescent signal from labeled SNAP-UbiquiNav-C942S vesicles are clearly visible along axons.

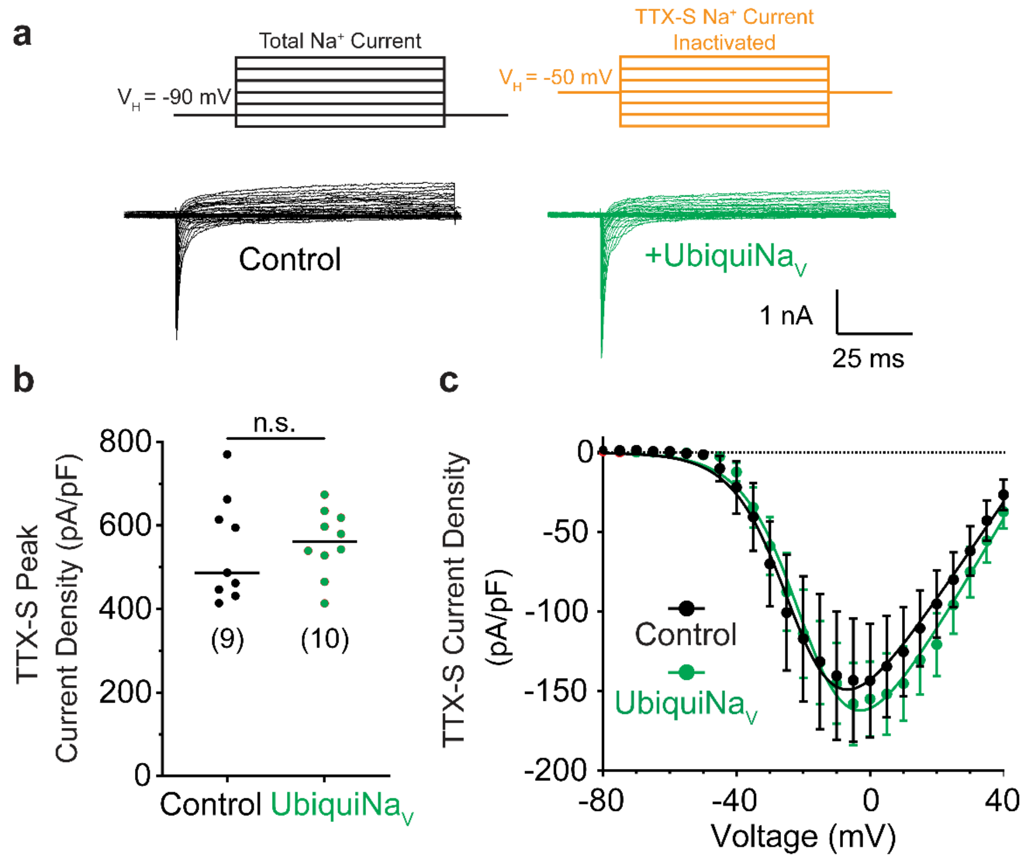


**Supplementary Figure 2. UbiquiNav does not affect Tetrodotoxin-sensitive Nav currents in rat pup DRG neurons**

(**a**) Pulse protocol used to isolate TTX-S currents in voltage-clamp recordings of 2-4d old rat pup DRG neurons. While holding at -90 mV, total $Na^+$ current was recorded. Then, the holding potential was changed to -50 to inactivate all TTX-S channels. By reference subtraction of the resulting currents, TTX-S current was quantified (rep traces shown). Neurons were transfected with either UbiquiNav (green traces) or eGFP-control (black traces).

(**b**) Peak inward current density of TTX-S currents from DRG neurons expressing either eGFP control (●; n=9) or UbiquiNav (●; n=10). n.s.- p>0.05 by Student's unpaired t-test.

(**c**) Current-voltage relationship of TTX-S currents recorded from DRG neurons expressing either eGFP control (●; n=9) or UbiquiNav (●; n=10).

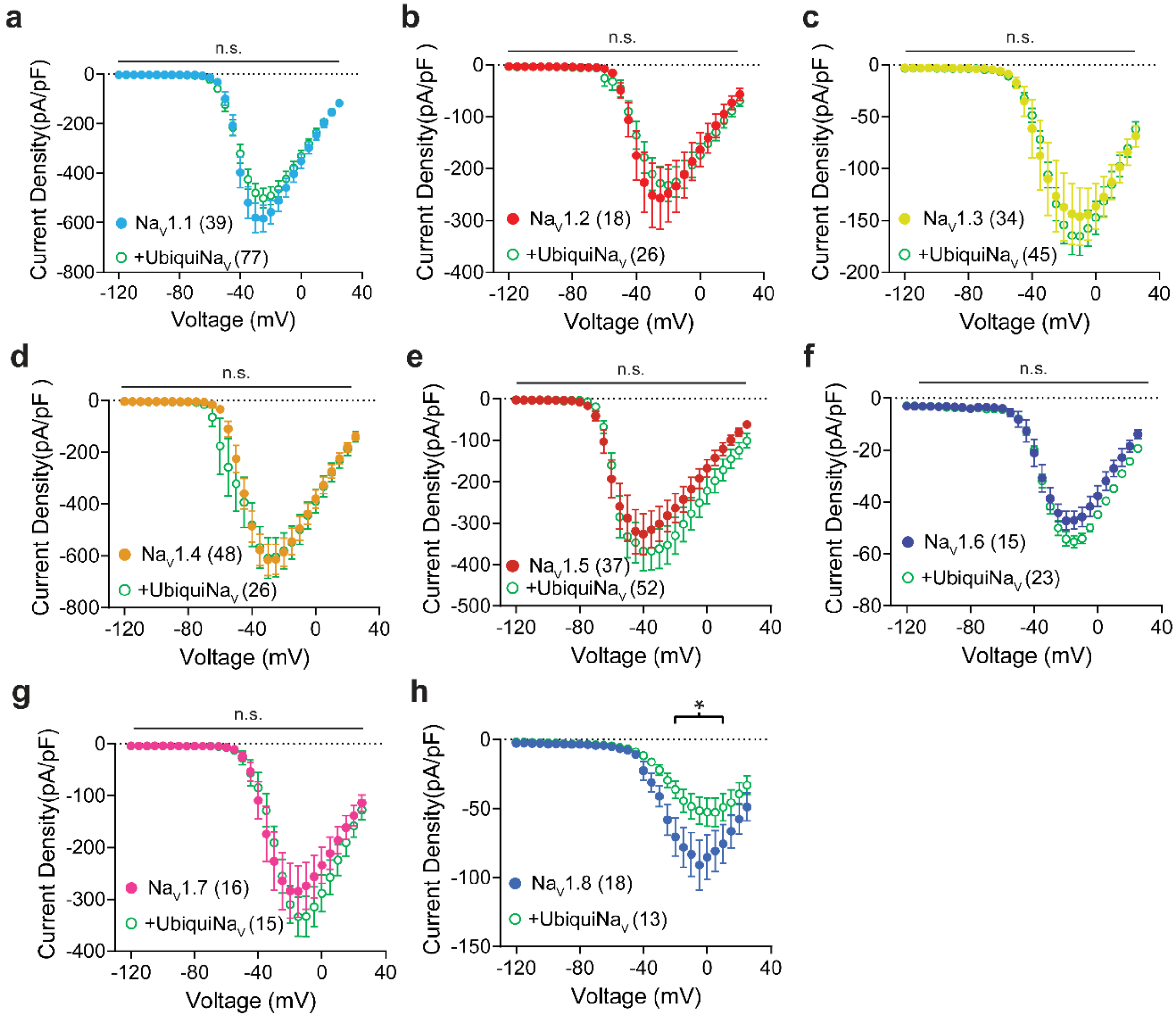


**Supplementary Figure 3. UbiquiNav is selective for Nav1.8 over other human Nav channels**

Current-voltage relationships from Expi293 cells expressing human Nav channel isoforms Nav1.1-Nav1.7 and UbiquiNav; DRG-derived ND7/23 cells were used to express hNav1.8. Cells were sorted by Flow Cytometry prior to whole cell voltage-clamp recording on the Sophion Qube 384 well Automated Patch Clamp system.

(**a**) Comparison of Nav1.1 current density in the presence of UbiquiNav (○) or mCherry control (●).

(**b**) Comparison of Nav1.2 current density in the presence of UbiquiNav (○) or mCherry control (●).

(**c**) Comparison of Nav1.3 current density in the presence of UbiquiNav (○) or mCherry control (●).

(**d**) Comparison of Nav1.4 current density in the presence of UbiquiNav (○) or mCherry control (●).

(**e**) Comparison of Nav1.5 current density in the presence of UbiquiNav (○) or mCherry control (●).

(**f**) Comparison of Nav1.6 current density in the presence of UbiquiNav (○) or mCherry control (●).

(**g**) Comparison of Nav1.7 current density in the presence of UbiquiNav (○) or mCherry control (●).

(**h**) Nav1.8 does not express well in HEK293 cells. ND7/23 cells were transfected with hNav1.8 prior to FACS and APC to enable comparison of Nav1.8 current density in the presence of UbiquiNav (○) or mCherry control (●).

n.s. $p>0.05$ by linear mixed-effects modeling.

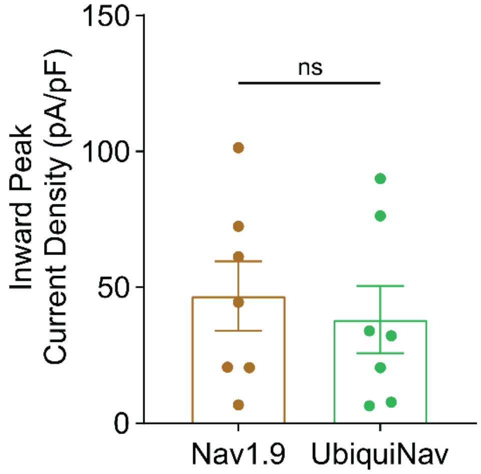


**Supplementary Figure 4. UbiquiNa$_V$ does not affect Na$_V$1.9 currents**

Peak inward current densities from manual whole-cell voltage clamp of rat pup superior cervical ganglion neurons expressing hNa$_V$1.9, mCherry, and either GFP control (●; n=7) or UbiquiNa$_V$-P2A-eGFP (●; n=7). n.s., $p > 0.05$ by Student's two-tailed t-test

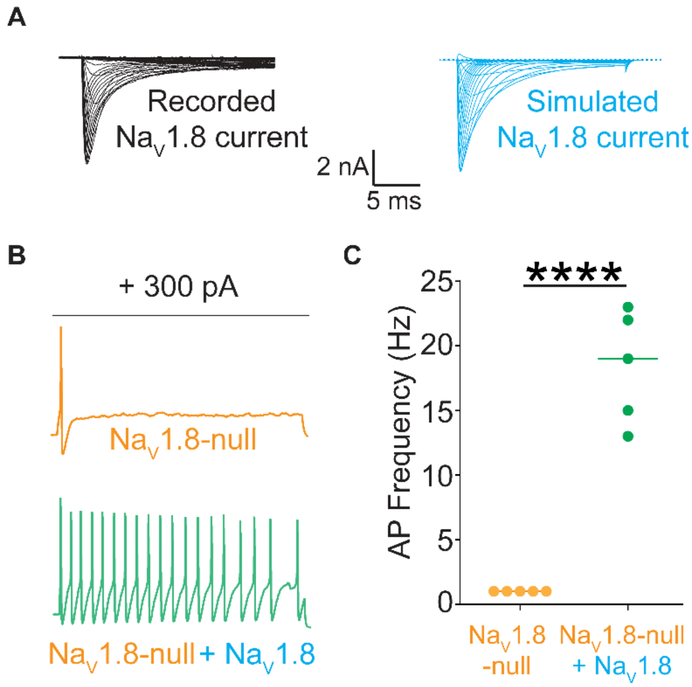


**Supplementary Figure 5. Dynamic clamp addition of Nav1.8 conductance restores repetitive firing in Nav1.8-null mouse DRG neurons**

(**a**) Dynamic clamp modeling (Hodgkin-Huxley) of Nav1.8 conductance yields simulated Nav1.8 currents (blue trace) that simulate recorded Nav1.8 currents (black trace) with high fidelity.

(**b**) Nav1.8-null mouse DRG neurons are incapable of repetitive firing (orange trace). Dynamic clamp addition of simulated Nav1.8 conductance restores repetitive firing capability in these neurons (green trace)

(**c**) Quantification of action potentials fired in response to a 300-pA stimulus before (●) and after (●) dynamic clamp addition of Nav1.8 conductance. **** $p<0.0001$ by Mann-Whitney U-test.

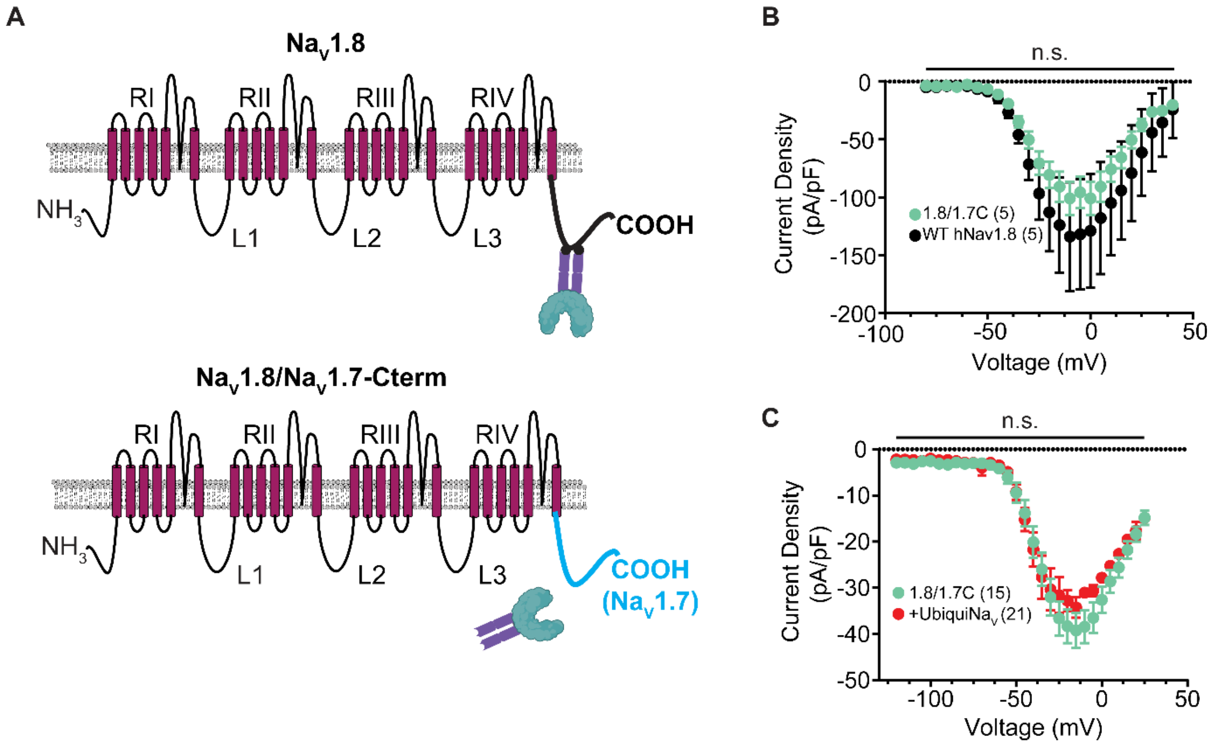


**Supplementary Figure 6. Replacing the Nav1.8 C-terminus with the Nav1.7 C-terminus yields a functional Nav1.8 channel resistant to UbiquiNav-mediated degradation.**

(**a**) Schematic of UbiquiNav binding to the Nav1.8 C-terminus. Swapping the C-terminus of Nav1.8 with the C-terminus of Nav1.7 removes the binding site of UbiquiNav from the Nav1.8 channel.

(**b**) Swapping the C-terminus of Nav1.8 with the C-terminus of Nav1.7 yields a channel that demonstrates a similar current-voltage relationship and expression profile with WT human Nav1.8.

(**c**) Nav1.8/1.7C can be expressed in Expi293 cells, unlike WT Nav1.8. Nav1.8/1.7C currents are not reduced in the presence of UbiquiNav (●) vs. mCherry control (●). n.s. $p>0.05$ by linear mixed effects modeling.
